# Cytokinin receptor AHK3 influences leaf size by modulating *trans*-zeatin-type cytokinin levels in xylem

**DOI:** 10.1101/2025.09.05.674380

**Authors:** Kota Monden, Takamasa Suzuki, Mikiko Kojima, Yumiko Takebayashi, Takehiro Kamiya, Takatoshi Kiba, Hitoshi Sakakibara, Tsuyoshi Nakagawa, Takushi Hachiya

## Abstract

*Trans*-zeatin (*t*Z)-type cytokinins (CKs) are synthesized in roots in response to nitrate, transported to shoot via xylem, and coordinate diverse physiological processes in aerial organs. Within this mechanism, the regulation of CK biosynthesis by nitrate signaling via NIN-like protein 7 as well as the loading of *t*Z-type CKs into xylem by ATP-binding cassette transporter G14 have been well studied. However, the roles of other components remain unclear. Here, we show that CK perception and degradation in roots, as mediated by Arabidopsis histidine kinase 3 (AHK3) and CK oxidase/dehydrogenase 4 (CKX4), modulate xylem *t*Z-type CKs transport and leaf CK status. Grafting experiments demonstrated that root-specific *AHK3* deficiency systemically increased leaf blade area through long-distance signals of root-derived *t*Z-type CK, perceived by shoot-expressed AHK3. Transcriptome and hormonome analyses revealed that root-specific *AHK3* deficiency reduced *CKX4* expression in roots, elevating *t*Z-type CK levels in roots and xylem sap and thereby enhancing leaf CK response. Transfer experiments manipulating root nitrate levels showed that root-specific *AHK3* deficiency promoted leaf blade area in a manner dependent on both nitrate and root-derived *t*Z-type CK signaling. Moreover, both nitrate signals and root-expressed AHK3 are required for maximal *CKX4* induction in roots, and root-specific *CKX4* deficiency enhanced leaf blade area in a nitrate-dependent manner. These findings reveal a novel mechanism in which an AHK3–CKX4 module governs xylem transport of *t*Z-type CKs, fine-tuning leaf size according to nitrogen availability in roots.

**SIGNIFICANCE STATEMENT:** To clarify a mechanism that attenuates *trans*-zeatin-type cytokinin transport from roots to shoots in response to nitrate signaling, this study examined the root-specific role of cytokinin receptors. Our results show that cytokinin perception and degradation, as mediated by Arabidopsis histidine kinase 3 and cytokinin oxidase/dehydrogenase 4, modulate xylem *trans*-zeatin-type cytokinin transport, thereby fine-tuning leaf cytokinin status and growth in a nitrate-dependent manner, providing new insights into long-distance cytokinin transport according to nitrogen availability.

## INTRODUCTION

Cytokinins (CKs) are essential plant hormones that coordinate diverse developmental and physiological processes, including cell division, shoot and root growth, vascular formation, and responses to stress and nutrients (Werner et al., 2001; Riefler et al., 2006; Matsumoto-Kitano et al., 2008; Sakakibara, 2021). In *Arabidopsis*, the biologically active CK forms include *N*^6^-(*Δ*^2^-isopentenyl) adenine (iP) and *trans*-zeatin (*t*Z). The initial and rate-limiting step of CK biosynthesis involves *N*^6^-prenylation of adenosine 5′-phosphates catalyzed by adenosine phosphate-isopentenyltransferases (IPTs), producing iP-riboside 5′-phosphates (iPRPs) (Takei et al., 2001a; Miyawaki et al., 2006). These intermediates are subsequently hydroxylated by cytochrome P450 monooxygenases CYP735A1 and CYP735A2 to form *t*Z-riboside 5′-phosphates (*t*ZRPs) (Takei et al., 2004a). Ribotide precursors are then converted into iP and *t*Z by CK-activating enzyme members of the lonely guy (LOG) family (Kurakawa et al., 2007; Kuroha et al., 2009). CK homeostasis is further modulated by reversible *O*-glucosylation and tentatively reversible *N*-glucosylation catalyzed by glucosyltransferases (UGTs), and by irreversible degradation by cytokinin oxidase/dehydrogenases (CKXs) (Werner et al., 2006; Wang et al., 2011; Jin et al., 2013; Wang et al., 2013; Hošek et al., 2020). CK perception and signaling rely on a multistep His-to-Asp phosphorelay system (Hutchison and Kieber, 2002; Müller and Sheen, 2007). Three membrane-spanning histidine kinases, Arabidopsis histidine kinase (AHK)2, AHK3, and CRE1/AHK4, act as CK receptors (Inoue et al., 2001). Upon ligand binding, the phosphoryl group is transferred from AHKs to histidine phosphotransfer proteins, which shuttle into the nucleus and activate type-B response regulators (Hutchison and Kieber, 2002; Müller and Sheen, 2007). These transcription factors induce CK-responsive genes, including type-A response regulators that function as negative feedback elements (To et al., 2004).

In addition to local roles, CKs act as long-distance signaling molecules that coordinate developmental and physiological processes between roots and shoots (Sakakibara, 2021). In *Arabidopsis*, *t*Z-type CKs predominate in xylem sap, whereas iP-type CKs are more abundant in phloem sap (Hirose et al., 2008; Osugi et al., 2017). This distribution reflects tissue-specific expression of CK biosynthetic genes: *CYP735A2* is strongly expressed in root vasculature, while *IPT3* is mainly expressed in phloem tissues of both roots and shoots (Miyawaki et al., 2004; Takei et al., 2004a,b; Kiba et al., 2013). These activities establish a directional flow of CK species, with root-derived *t*Z-type CKs transported to shoots via xylem and shoot-derived iP-type CKs transported to roots via phloem tissue (Hirose et al., 2008; Matsumoto-Kitano et al., 2008; Kiba et al., 2013). A central component of the root-to-shoot CK transport system is ATP-binding cassette transporter G14 (ABCG14), which loads CKs into xylem in root stele and also contributes to phloem unloading in shoots (Ko et al., 2014; Zhang et al., 2014; Zhao et al., 2021). Importantly, root-to-shoot transport of *t*Z-type CKs mediated by ABCG14 is essential for maintaining shoot growth, which is supported by the observation that *abcg14* mutants show strongly reduced shoot growth during both the vegetative and reproductive stages (Ko et al., 2014; Zhang et al., 2014). In addition, heterologous transport assays in yeast have shown that ABCG14 transports multiple CK species, including both iP- and *t*Z-types, underscoring its pivotal role in systemic CK distribution and signaling (Zhao et al., 2023).

Root-to-shoot *t*Z-type CK transport is strongly influenced by plant nitrogen status. In *Arabidopsis*, nitrate supply induces the expression of *IPT3* and *CYP735A2* in roots and increases the concentrations of *t*ZR and *t*ZRMP (Takei et al., 2004b; Maeda et al., 2018). Nitrate supply also enhances the xylem transport of *t*Z-riboside (*t*ZR) and activation of CK signaling in leaves (Sakakibara et al., 1998; Takei et al., 2001b; Osugi et al., 2017; Abualia et al., 2022). Furthermore, NIN-like protein 7 (NLP7), a nitrate sensor, drives the nitrate-dependent induction of *IPT3* and *CYP735A2* (Liu et al., 2022). Therefore, nitrate-induced increases in *t*Z and *t*ZR concentrations and in *ARR5* promoter activity in shoots are markedly reduced in *nlp7* mutants (Abualia et al., 2022). Nitrate signaling also induces *ABCG14* expression in roots in an NLP7-dependent manner (Abualia et al., 2022). In addition, long-distance signals of root-derived *t*Z and *t*ZR differentially affect leaf growth traits, and their ratios in xylem sap are known to be modulated by nitrate supply (Osugi et al., 2017). Taken together, these findings indicate that plants adjust leaf growth according to root nitrate status mediated by *t*Z-type CKs, and that NLP7 and ABCG14 act as key components governing the root-to-shoot transport of *t*Z-type CKs. On the other hand, the feedback mechanism that attenuates the nitrate-induced accumulation of *t*Z-type CKs remains poorly understood. The nitrate-inducible GARP-type transcriptional repressor 1s (NIGT1s) are induced by NLPs and repress *IPT3* and *CYP735A2* expression, suggesting that they may function as feedback components (Kiba et al., 2018; Maeda et al., 2018). However, because *nigt1.1/1.2/1.3/1.4* quadruple mutants show only a modest increase in *t*Z-type CK concentrations (Maeda et al., 2018), other major feedback components that modulate *t*Z-type CK levels likely remain unidentified.

One promising candidate for a key feedback component may be AHK3. In *Arabidopsis*, disruption of *AHK3* causes a pronounced twofold to threefold increase in all *t*Z-type CK species (Riefler et al., 2006). Moreover, our previous study proposed that root-specific deficiencies of *AHK2* and *AHK3* increased *t*Z-type CKs concentration in roots, which in turn affected *t*Z-type CKs concentrations, type-A *ARRs* expressions, and growth in the shoot (Monden et al., 2022). Furthermore, *AHK3* was identified as one of the top genes co-expressed with *NLP7* (Supplementary Table S1), suggesting that AHK3 may act in association with nitrate responses via NLP7. In addition, AHK3 has been reported to exhibit a higher affinity for *t*Z than for iP (Stolz et al., 2011), implying a preferential role in *t*Z-type CK signaling. Together, these findings suggest that root-expressed AHK3 may modulate root-to-shoot transport of *t*Z-type CKs in response to nitrogen availability in roots, although the underlying mechanism and physiological relevance remain unclear.

In this study, we show that CK perception and degradation in roots, mediated by AHK3 and CKX4, modulate *t*Z-type CK levels in the xylem, thereby influencing CK responses and growth in leaves. Growth analyses of grafted plants demonstrated that root-specific *AHK3* deficiency enlarged leaf blade area through long-distance signals of root-derived *t*Z-type CK, perceived by shoot-expressed AHK3. Transcriptome and hormonome analysis further revealed that root-specific *AHK3* deficiency reduced *CKX4* expression in roots, elevating *t*Z-type CK levels in roots and xylem sap and thereby enhancing CK responses in leaves. Transfer experiments manipulating root nitrate levels showed that nitrate-induced leaf growth promotion depends on long-distance signals of root-derived *t*Z-type CK, which is in turn controlled by root-expressed AHK3 and CKX4. We propose a mechanism in which the AHK3–CKX4 module governs xylem transport of *t*Z-type CK in response to nitrogen availability in roots, thereby fine-tuning leaf size to maintain balanced plant growth.

## RESULTS

### Root-expressed AHK3 modulates shoot and root growth

To identify which root-expressed CK receptor(s) influence shoot and root growth, we grafted Col-0 (WT) scions onto rootstocks derived from WT, single mutant (i.e., *ahk2-5*; *ahk2*, *ahk3-7*; *ahk3*, *cre1-2*; *ahk4*) or double mutant (i.e., *ahk2-5 ahk3-7*; *ahk23*, *ahk2-5 cre1-2*; *ahk24*, *ahk3-7 cre1-2*; *ahk34*) five-day-old seedlings (Riefler et al., 2006). Five days after grafting, we selected plants (scion/rootstock: WT/WT, WT/*ahk2*, WT/*ahk3*, WT/*ahk4*, WT/*ahk23*, WT/*ahk24*, WT/*ahk34*) for further analysis if they showed vigorous growth without adventitious roots. Grafted plants were then cultivated for 7 days on either half-strength MS (1/2 MS) medium or nutrient-rich soil. We measured shoot fresh weight, leaf blade area, and leaf number. Overall, WT/*ahk3*, WT/*ahk23*, and WT/*ahk34* displayed greater shoot growth than WT/WT (Fig. 1A, Supplementary Fig. S1A). Relative to the WT/WT control, these lines showed an increase in shoot fresh weight of 26%, 35%, and 19%, and in leaf blade area of 26%, 32%, and 17%, respectively (Fig. 1, B and C). Under nutrient-rich soil conditions, shoot fresh weight increased by 34%, 25%, and 22%, and leaf blade area by 32%, 21%, and 17% in WT/*ahk3*, WT/*ahk23*, and WT/*ahk34*, respectively, compared with WT/WT (Supplementary Fig. S1, B and C).

**Figure 1.**
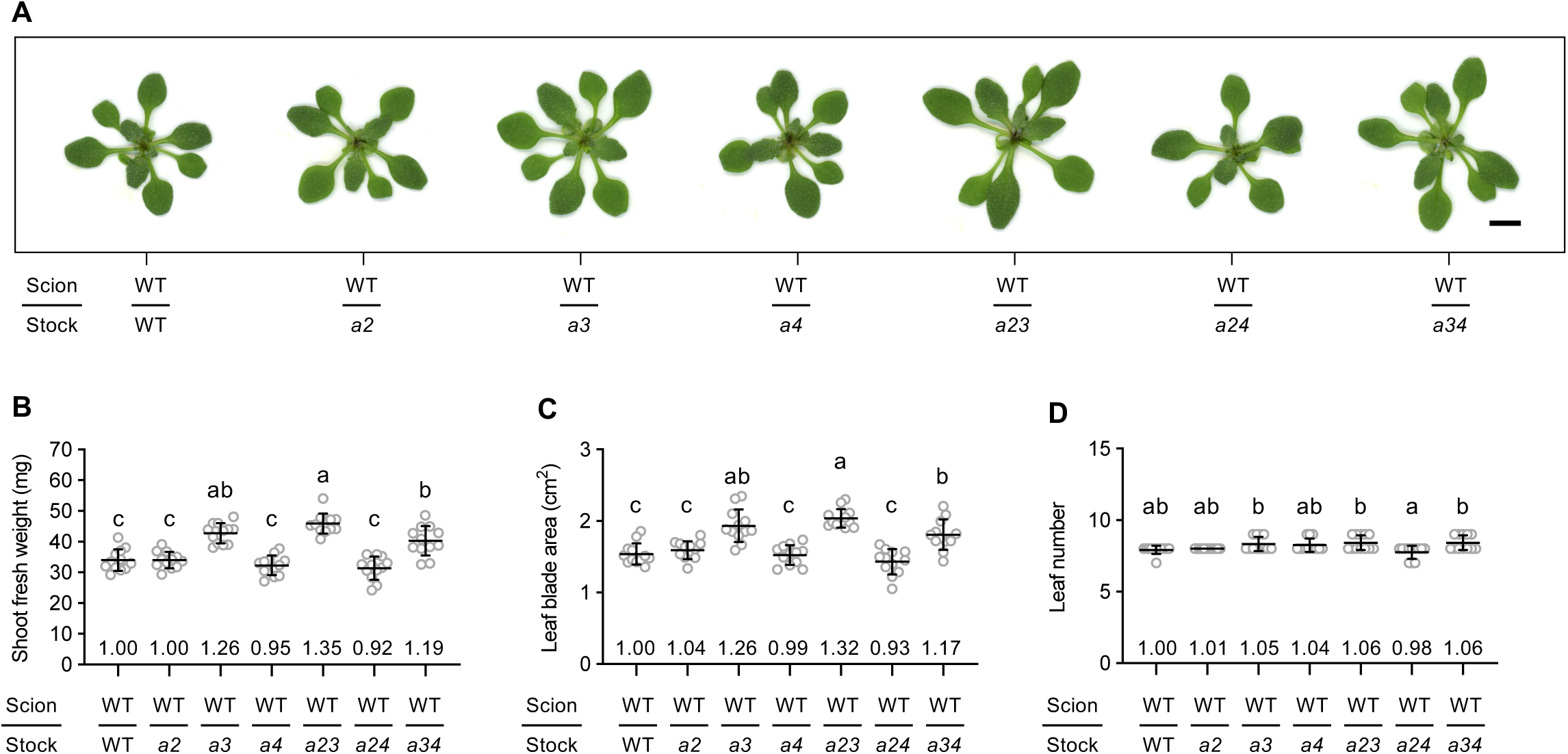
Effects of root-specific CK receptor deficiency on shoot growth in medium. (A) Representative shoots of grafted plants. Scale bars = 5 mm. (B–D) Shoot fresh weight (B), leaf blade area (C), and leaf number (D) of grafted plants. Data pooled from two independent grafting experiments are shown as mean ± SD (n = 12). Different lowercase letters indicate significant differences, as determined via Tukey–Kramer test (*P* < 0.05). Numbers on graphs show mean values relative to WT/WT (set = 1). Grafted plants are denoted as “scion line/rootstock line.” WT, Col-0; *a2*, *ahk2-5*; *a3*, *ahk3-7*; *a4*, *cre1-2*; *a23*, *ahk2-5 ahk3-7*; *a24*, *ahk2-5 cre1-2*; *a34*, *ahk3-7 cre1-2*.

Leaf number was generally similar across grafted plants in both conditions (Fig. 1D, Supplementary Fig. S1D). We next monitored primary root length in grafted plants transferred to 1/2 MS medium over a five-day period. Primary root length was comparable at day 0, but WT/*ahk3*, WT/*ahk23*, and WT/*ahk34* developed significantly shorter roots than WT/WT from day 3 onward (Supplementary Fig. S2). In contrast, WT/*ahk2*, WT/*ahk4*, and WT/*ahk24* showed no significant differences in shoot or root growth relative to WT/WT (Fig. 1, Supplementary Figs. S1 and S2). Notably, the relative growth of WT/*ahk3* was comparable to WT/*ahk23* and WT/*ahk34*, indicating that loss of AHK3 alone is sufficient to alter overall plant growth. We therefore focused subsequent analyses on WT/*ahk3*, using WT/WT, WT/*ahk2*, and WT/*ahk4* as comparisons.

### Root-expressed AHK3 modulates root CK levels

Our previous work showed that root-specific deficiencies of *AHK2* and *AHK3* markedly increased iP- and *t*Z-type CK levels in roots, but the receptor responsible for governing root CK levels was unclear (Monden et al., 2022). To clarify this, we measured iP- and *t*Z-type CK concentrations in WT/WT, WT/*ahk2*, WT/*ahk3*, and WT/*ahk4* roots. Root concentrations of nearly all iP- and *t*Z-type CK species were significantly elevated in WT/*ahk3* compared with WT/WT (Fig. 2, Supplementary Fig. S3). In contrast, root iP-and *t*Z-type CKs were largely unchanged in WT/*ahk2* and WT/*ahk4* relative to WT/WT, except for iP7G and iP9G in WT/*ahk4* (Fig. 2, Supplementary Fig. S3). These findings indicate that AHK3 expressed in roots modulates CK homeostasis.

**Figure 2.**
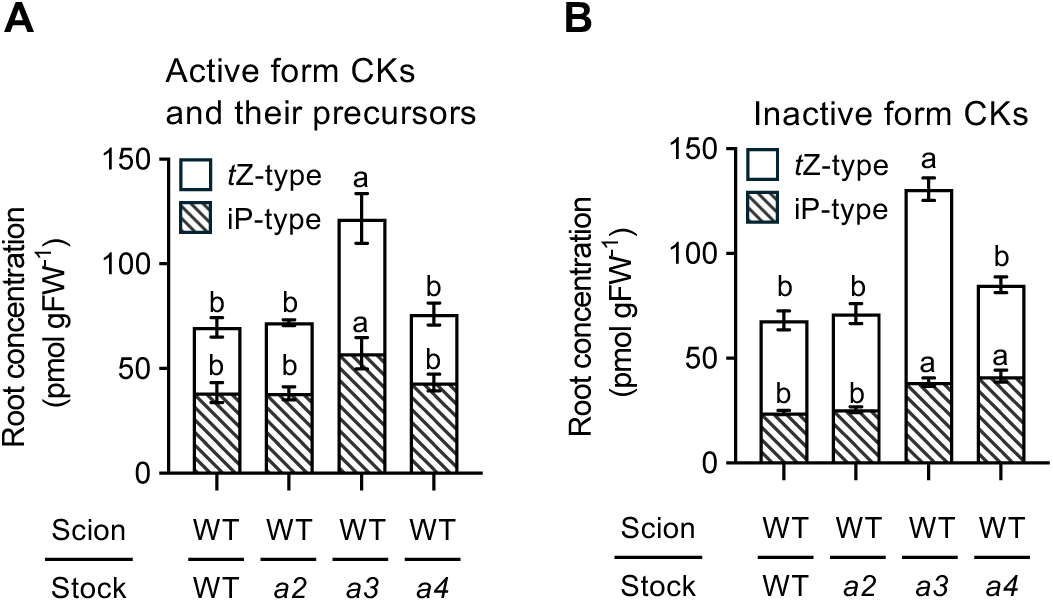
Effects of root-specific CK receptor deficiency on root CK concentration. (A–B) Root concentrations of active form iP and *t*Z and their precursors (A), and inactive form iP- and *t*Z-type CKs (B) in grafted plants. Data are pooled from three independent grafting experiments and are reported as mean ± SD (n = 3). Different lowercase letters indicate significant differences, as determined via Tukey–Kramer test (*P* < 0.05). Grafted plants are denoted as “scion line/rootstock line.” WT, Col-0; *a2*, *ahk2-5*; *a3*, *ahk3-7*; *a4*, *cre1-2*; FW, fresh weight; *t*Z, *trans*-zeatin; iP, *N*^6^-(*Δ*^2^-isopentenyl) adenine; iP and their precursors = sum of iP, iPR, and iPRPs; inactive form iP-type CKs = sum of iP7G and iP9G; *t*Z and their precursors = sum of *t*Z, *t*ZR, and *t*ZRPs; inactive form *t*Z-type CKs = sum of *t*Z7G, *t*Z9G, *t*ZOG, and *t*ZROG.

### Root-expressed AHK3 influences the root transcriptome

Despite the fact that *AHK2*, *AHK3*, and *AHK4* are all expressed in roots (Higuchi et al., 2004), only AHK3 strongly affected root CK levels. This suggests that each CK receptor may regulate a distinct set of downstream genes. To test this hypothesis, we performed an RNA-seq analysis of root samples taken from WT/WT, WT/*ahk2*, WT/*ahk3*, and WT/*ahk4*. All data, including raw read counts, reads per million mapped reads (RPM), and normalized transcript levels are presented in Supplementary Tables S2–4. First, we examined differentially expressed genes (DEGs) between the roots of the WT/WT control and three grafted plants, i.e., WT/*ahk2*, WT/*ahk3*, and WT/*ahk4* (Supplementary Table S6). An analysis of the DEGs with a fold change of 1.5 or greater revealed that 62 genes were significantly upregulated in WT/*ahk3* relative to WT/WT roots, whereas 91 genes were downregulated (Fig. 3A, Supplementary Figs. S4 and S5). Notably, relatively few DEGs were detectable in the roots of WT/*ahk2* and WT/*ahk4* (Fig. 3A, Supplementary Figs. S4 and S5). Next, we conducted a Gene Ontology (GO) term enrichment analysis of the 62 upregulated genes and the 87 downregulated genes (excluding *AHK3*) identified in WT/*ahk3* roots. This revealed that multiple hormone-related terms, such as “Cellular response to hormone stimulus” and “Hormone catabolic process,” were enriched in both sets (Supplementary Fig. S6). To further examine the extent to which CK receptor regulates CK signaling-related genes in roots, we analyzed the expression of CK-inducible and -repressible genes in the roots of grafted plants; to do so we used a curated gene list (Bhargava et al., 2013). We found that the expression levels of both CK-inducible and -repressible genes in the roots of WT/*ahk4* were comparable to the levels detected in WT/WT (Fig. 3B). In contrast, the expression of CK-inducible genes in root tissue samples was significantly lower in both WT/*ahk2* and WT/*ahk3* relative to the WT/WT (Fig. 3B). Moreover, CK-repressible genes were significantly increased in WT/*ahk3* and showed an increasing trend in WT/*ahk2* (Fig. 3B). Notably, the magnitude of these changes was more pronounced in WT/*ahk3* than in WT/*ahk2* roots, suggesting that root-expressed AHK3 plays a major role in regulating root CK signaling.

**Figure 3.**
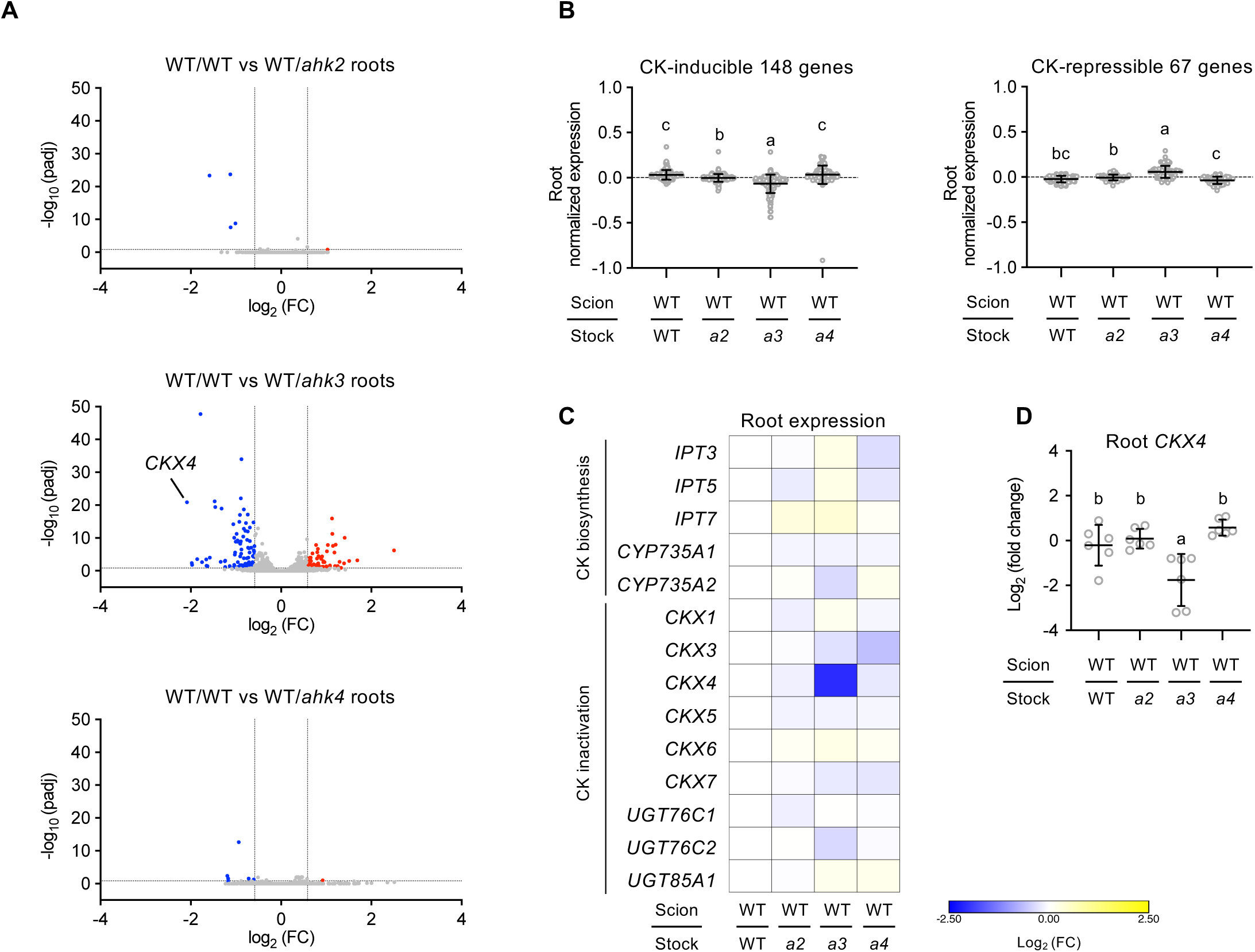
Effects of root-specific CK receptor deficiency on the root transcriptome. (A) Volcano plots of DEGs between WT/WT and WT/*ahk2*, WT/*ahk3*, or WT/*ahk4* roots. DEG analysis was performed with RNAseqChef (Etoh et al., 2023) using edgeR, applying a 1.5-fold change threshold and FDR cutoff of 0.15 (all other parameters default). x-axis: log_2_ fold change (log_2_ FC); y-axis: –log_10_ adjusted p-value (padj). Each dot represents a single gene. Red, blue, and grey indicate upregulated, downregulated, and non-significant genes, respectively. Log_2_ FC was limited to –4 to 4 and padj to ≦50. Full-range volcano plots are provided in Supplementary Fig. S4. (B) Mean normalized transcript levels of CK-inducible and CK-repressible genes (mean ± SD). Gene list obtained from Bhargava *et al*. (2013). (C) Transcriptional changes in CK biosynthesis and inactivation genes, visualized as heatmaps with MORPHEUS (https://software.broadinstitute.org/morpheus) based on log_2_-transformed ratios of each graft to WT/WT mean RPM. Color scale: blue (minimum) to yellow (maximum) per row. (D) Root expression of *CKX4* in grafted plants by qRT-PCR. Relative transcript levels were normalized to the WT/WT root mean (set = 1) and log_2_-transformed. Data pooled from two independent grafting experiments are shown as mean ± SD (n = 6). Different lowercase letters indicate significant differences, as determined via Tukey–Kramer test (*P* < 0.05). Grafted plants are denoted as “scion line/rootstock line.” WT, Col-0; *a2*, *ahk2-5*; *a3*, *ahk3-7*; *a4*, *cre1-2*; FC, fold change.

Next, to identify candidate gene(s) that may be responsible for increased CK levels in WT/*ahk3* roots, we examined whether the DEGs observed in WT/*ahk3* roots included genes that are known to be involved in CK biosynthesis and inactivation. Of these genes, only *CKX4*, which is involved in CK degradation, was among the genes that were downregulated genes in WT/*ahk3* roots relative to the WT/WT (Fig. 3, A and C). Quantitative RT-PCR (qRT-PCR) analysis confirmed that expression of *CKX4* in roots was significantly decreased in WT/*ahk3* relative to WT/WT (Fig. 3D). Taken together, these results suggest that root-expressed AHK3 modulates root CK levels, potentially via CKX4-mediated CK degradation.

### Root-expressed AHK3 modulates *t*Z-type CKs in xylem sap

CKs synthesized in roots are transported to shoots mainly via xylem pathway (Ko et al., 2014; Osugi et al., 2017). Given that we observed relatively high CK levels in WT/*ahk3* roots, we hypothesized that the CK levels of xylem sap may also be altered. To verify this, we measured the CK concentrations of xylem sap samples from WT/WT, WT/*ahk2*, WT/*ahk3*, and WT/*ahk4*. We found that *t*Z, *t*ZR, and *t*ZRPs were relatively more abundant than inactive forms of *t*Z-type CK and iP-type CKs in WT/WT xylem sap (Supplementary Fig. S7A). This suggests that there is preferential root-to-shoot transport of *t*Z and its precursors. Notably, the concentrations of *t*Z-type CKs were significantly higher in WT/*ahk3* than in WT/WT xylem sap, while the concentrations of iP-type CKs remained largely unchanged (Fig. 4, Supplementary Fig. S7B). Among the major CK species detected in xylem sap, the concentration of *t*Z showed a 2.22-fold increase in WT/*ahk3* relative to WT/WT xylem sap (Fig. 4B). In contrast, the concentrations of *t*ZR and *t*ZRPs were both slightly higher in WT/*ahk3* than in WT/WT xylem sap, showing 1.20-fold and 1.37-fold increases, respectively (Fig. 4B). Finally, the concentrations of iP- and *t*Z-type CKs were largely unchanged in WT/*ahk2* and WT/*ahk4* relative to WT/WT xylem sap, except for *t*Z and *t*ZRPs in WT/*ahk4* (Fig. 4, Supplementary Fig. S7B). Overall, these results suggest that root-expressed AHK3 modulates the xylem transport of *t*Z-type CKs, especially *t*Z.

**Figure 4.**
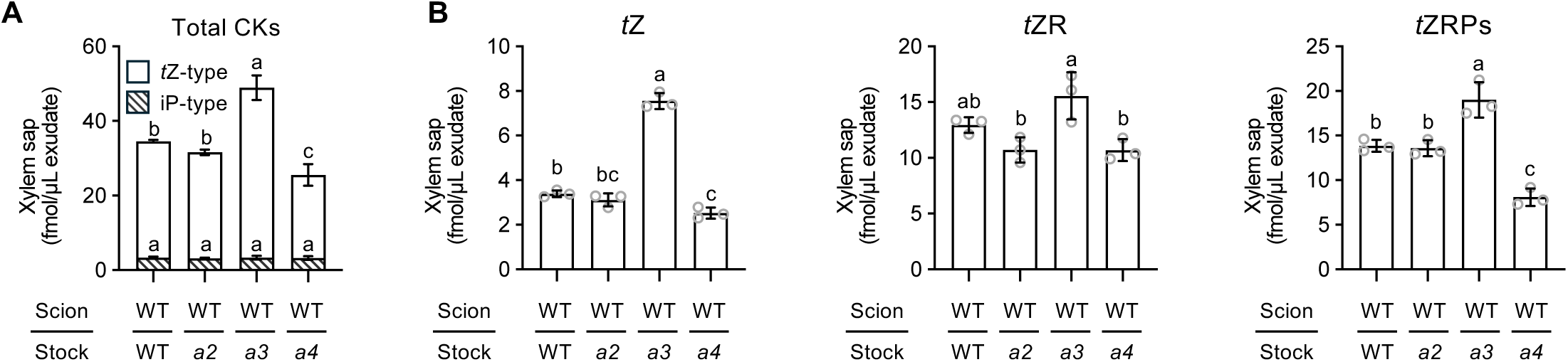
Effects of root-specific CK receptor deficiency on CK concentration of xylem sap. (A–B) Xylem sap concentrations of iP- and *t*Z-type CKs (A) and of *t*Z, *t*ZR, and *t*ZRPs (B) in grafted plants. Data pooled from three independent grafting experiments are reported as mean ± SD (n = 3). Different lowercase letters indicate significant differences, as determined via Tukey–Kramer test (*P* < 0.05). Grafted plants are denoted as “scion line/rootstock line.” WT, Col-0; *a2*, *ahk2-5*; *a3*, *ahk3-7*; *a4*, *cre1-2*; *t*Z, *trans*-zeatin; *t*ZR, *t*Z-riboside; *t*ZRPs, *t*ZR 5′-phosphates; iP-type CKs = sum of iPR, iPRPs, and iP7G; *t*Z-type CKs = sum of *t*Z, *t*ZR, *t*ZRPs, *t*Z7G, *t*Z9G, and *t*ZOG.

### Root-expressed AHK3 systemically influences the shoot transcriptome

Next, to test whether elevated *t*Z-type CK levels in WT/*ahk3* xylem sap affected shoot CK levels, we measured CK concentrations in WT/WT, WT/*ahk2*, WT/*ahk3*, and WT/*ahk4* shoots. Unexpectedly, *t*Z-type CK concentrations were not higher in WT/*ahk3* than in WT/WT shoots, except for *t*Z9G (Supplementary Fig. S8). Similarly, almost all CK species in WT/*ahk2* and WT/*ahk4* showed no marked differences compared with WT/WT (Supplementary Fig. S8). We next analyzed whether root-specific CK receptor deficiency may alter the shoot transcriptomes. To do so, we performed RNA-seq on WT/WT, WT/*ahk2*, WT/*ahk3*, and WT/*ahk4* shoots. Raw read counts, RPM, and normalized transcript levels are provided in Supplementary Tables S2, S3, and S5. Differential expression analysis with a 1.5-fold cutoff revealed 163 genes significantly upregulated and 35 genes downregulated in WT/*ahk3* relative to WT/WT shoots (Fig. 5A, Supplementary Table S7). No DEGs were detected in WT/*ahk2* or WT/*ahk4* shoots (Fig. 5A, Supplementary Table S7). Moreover, the number of upregulated genes was about five times greater than the number of downregulated genes in WT/*ahk3* shoots (Fig. 5A, Supplementary Table S7). This finding indicates that root-specific *AHK3* deficiency mainly induces gene expression in shoots.

**Figure 5.**
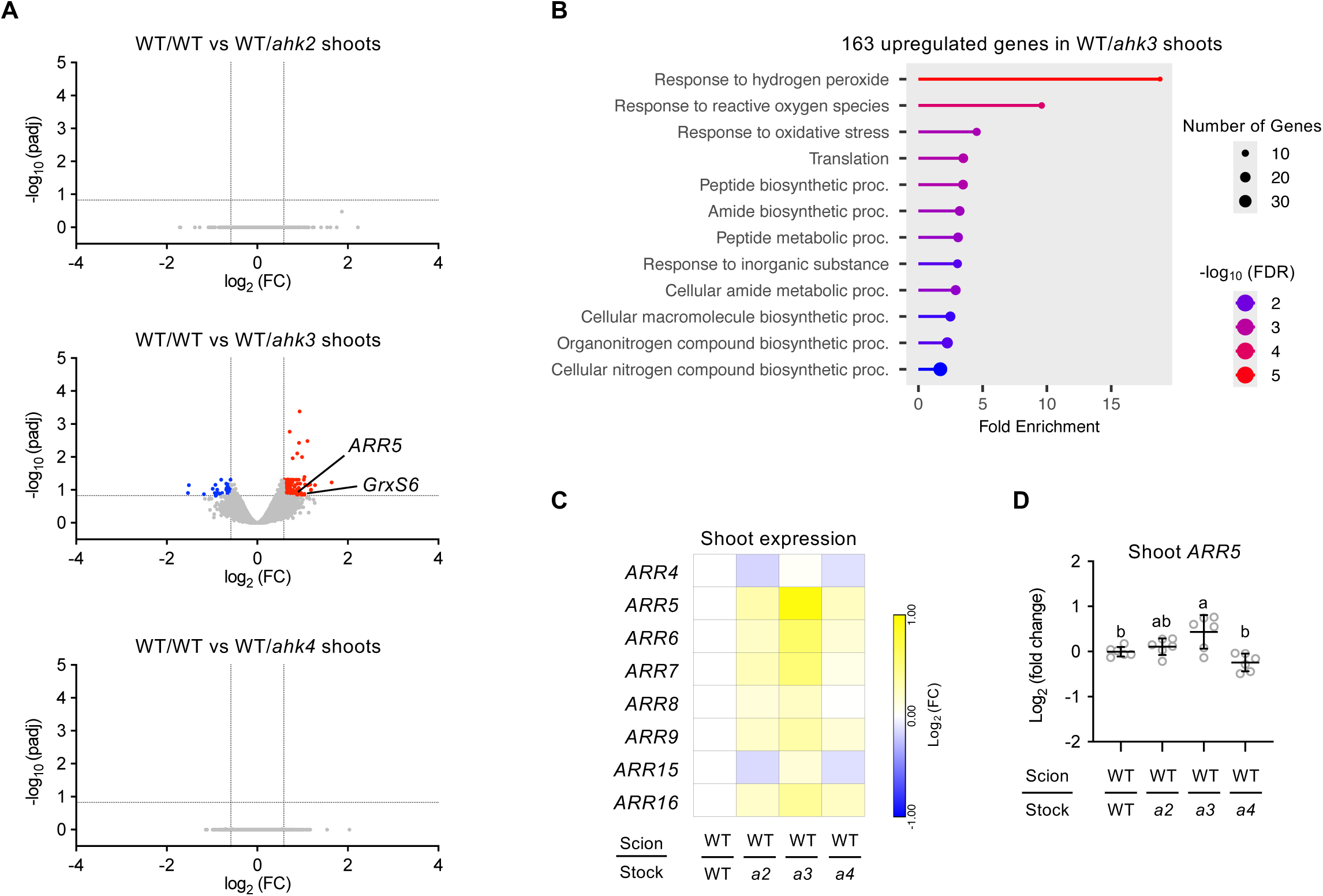
Effects of root-specific CK receptor deficiency on the shoot transcriptome. (A) Volcano plots of DEGs between WT/WT and WT/*ahk2*, WT/*ahk3*, or WT/*ahk4* shoots. DEG analysis was performed with RNAseqChef (Etoh et al., 2023) using edgeR, applying a 1.5-fold change threshold and FDR cutoff of 0.15 (default parameters otherwise). x-axis: log_2_ fold change (log_2_ FC); y-axis: –log_10_ adjusted p-value (padj). Each dot represents one gene; red, blue, and grey indicate upregulated, downregulated, and non-significant genes, respectively. Log_2_ FC values were limited to –4 to 4 and padj values to ≦5. (B) Lollipop plots showing GO enrichment analysis from the 163 genes upregulated in WT/*ahk3* shoots, generated with ShinyGO (Ge et al., 2020) using default settings for biological process terms. (C) Heatmaps of transcriptional changes in type-A *ARR* genes, visualized with MORPHEUS (https://software.broadinstitute.org/morpheus) based on log_2_-transformed ratios of each graft to the WT/WT mean RPM. Color scale: blue (minimum) to yellow (maximum) per row. (D) Shoot expression of *ARR5* determined by qRT-PCR. Values were normalized to the WT/WT shoot mean (set = 1) and log_2_-transformed. Data pooled from two independent grafting experiments are reported as mean ± SD (n = 6). Different lowercase letters indicate significant differences, as determined via Tukey–Kramer test (*P* < 0.05). Grafted plants are denoted as “scion line/rootstock line.” WT, Col-0; *a2*, *ahk2-5*; *a3*, *ahk3-7*; *a4*, *cre1-2*; FC, fold change.

To gain insight into the functional characteristics of the upregulated set, we performed GO enrichment analysis using the set of 163 upregulated genes. Enriched categories included “Translation,” “Response to inorganic substance,” and “Cellular nitrogen compound biosynthetic process” (Fig. 5B), indicating activation of genes related to protein synthesis, nutrient responses, and nitrogen metabolism in WT/*ahk3* shoots. We then examined whether these transcriptional changes correlated with altered shoot element concentrations. We found that the shoot concentrations of calcium, sulfur, manganese, and molybdenum were significantly higher in WT/*ahk3* than in WT/WT (Supplementary Fig. S9), suggesting a possible association between these nutrients and altered gene expression in WT/*ahk3* shoots. In contrast, although several nitrogen-related GO terms were enriched, shoot concentrations of nitrogen and nitrate did not differ between WT/WT and WT/*ahk3* (Supplementary Fig. S9). Thus, nitrogen status in WT/*ahk3* shoots did not drive the transcriptional changes observed in protein synthesis and nitrogen metabolism genes. Because nitrogen is a major protein component and both nitrate and CK signaling promote expressions of protein synthesis-related genes (Sakakibara et al., 2006), altered CK response in WT/*ahk3* shoots may underlie these transcriptional effects.

To further assess CK response, we compared the 163 upregulated genes in WT/*ahk3* shoots with CK-inducible genes (Bhargava et al., 2013). *Arabidopsis response regulator 5* (*ARR5*) and *Glutaredoxin-S6* (*GrxS6*) were among the overlapping genes (Fig. 5A). *ARR5*, one of ten type-A *ARRs*, is a representative CK-responsive gene functioning as a negative feedback regulator of CK signaling (To et al., 2004). Expression of several other type-A *ARRs* also tended to increase in WT/*ahk3* relative to WT/WT (Fig. 5C). Consistent with the RNA-seq results, qRT-PCR confirmed that *ARR5* expression was significantly higher in WT/*ahk3* shoots than in WT/WT (Fig. 5D). Taken together, these results indicate that root-expressed AHK3 modulates shoot gene expression, at least partly through activation of CK signaling.

### Root-derived CK signals are required for enhanced shoot growth via root-expressed AHK3

Elevated *t*Z-type CKs in WT/*ahk3* xylem sap suggested that long-distance signals of root-derived *t*Z-type CK contribute to enhanced shoot growth. To test this, we generated *ahk3 abcg14* and *ahk3 cyp735a1 cyp735a2* (*ahk3 cypDM*) mutants by traditional crossing (Supplementary Fig. S10A). Since both *abcg14* and *cypDM* shoots display lack proper root-derived *t*Z-type CK signaling (Kiba et al., 2013; Ko et al., 2014), these mutants allow assessment of its contribution. Disruption of the respective genes in *ahk3 abcg14* and *ahk3 cypDM* was confirmed by RT-PCR (Supplementary Fig. S10B). Grafted plants (scion/rootstock: WT/WT, WT/*ahk3*, WT/*abcg14*, WT/*ahk3 abcg14*, WT/*cypDM*, WT/*ahk3 cypDM*) were cultivated on 1/2 MS medium, and shoot growth was measured. Shoot fresh weight and leaf blade area were significantly reduced in WT/*abcg14* and WT/*ahk3 abcg14*, while WT/*cypDM* and WT/*ahk3 cypDM* showed no decrease compared with WT/WT (Fig. 6, A and B). These results agree with previous findings that root-specific deficiency of *ABCG14* impairs shoot growth, whereas deficiencies of *CYP735A1* and *CYP735A2* do not (Kiba et al., 2013; Ko et al., 2014). Notably, the increases in shoot fresh weight and leaf blade area seen in WT/*ahk3* compared with WT/WT were largely suppressed by disrupting *ABCG14* and completely abolished by disrupting *CYP735A1* and *CYP735A2* in roots (Fig. 6, A and B). These findings indicate that long-distance signals of root-derived *t*Z-type CKs are required for enhanced shoot growth in WT/*ahk3*.

**Figure 6.**
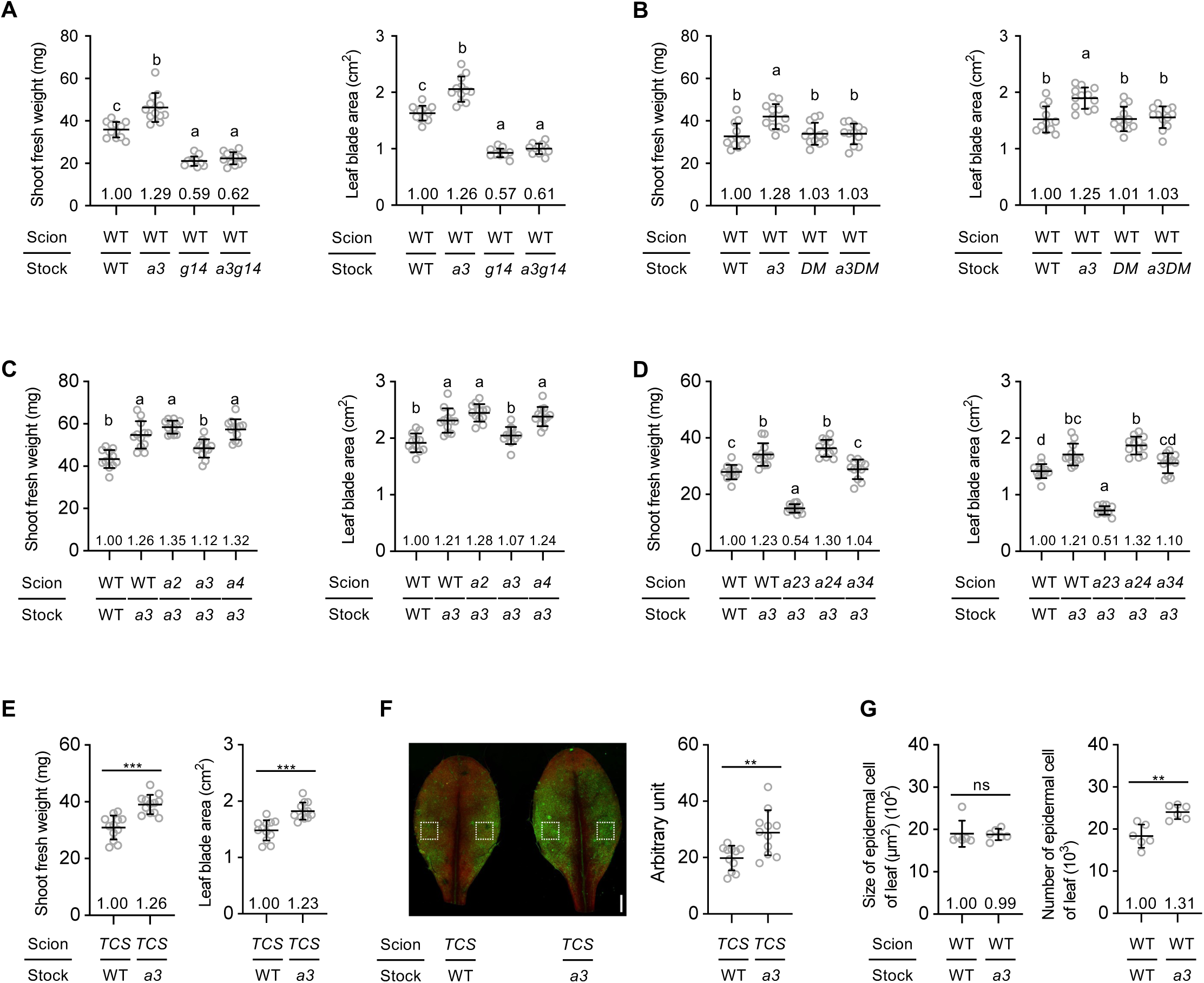
Effects of root-specific *AHK3* deficiency on shoot growth via CK signaling. (A–E) Shoot fresh weight and leaf blade area in grafted plants: (A) WT/WT, WT/*ahk3*, WT/*abcg14*, and WT/*ahk3 abcg14*; (B) WT/WT, WT/*ahk3*, WT/*cypDM*, and WT/*ahk3 cypDM*; (C) WT/WT, WT/*ahk3*, *ahk2*/*ahk3*, *ahk3*/*ahk3*, and *ahk4*/*ahk3*; (D) WT/WT, WT/*ahk3*, *ahk23*/*ahk3*, *ahk24*/*ahk3*, and *ahk34*/*ahk3*; (E) *TCSn:GFP*/WT and *TCSn:GFP*/*ahk3*. (F) Representative fluorescent image of the adaxial side of the sixth true leaf in grafted plants. Green = GFP; red = autofluorescence. Scale bars = 1 mm. GFP intensity was quantified with ImageJ as the mean signal in the region marked by a white dashed box. (G) Epidermal cell size and number on the abaxial side of the fifth true leaf. Data pooled from two independent grafting experiments are reported as mean ± SD (n = 12 (A–E), 11 (F), or 6 (G)). Different lowercase letters indicate significant differences, as determined via Tukey–Kramer test (*P* < 0.05). Asterisks indicate significant differences (\*\**P* < 0.01, \*\*\**P* < 0.001, Welch’s *t*-test). Graph numbers show mean values relative to WT/WT (set = 1). Grafted plants are denoted as “scion line/rootstock line.” WT, Col-0; *a2*, *ahk2-5*; *a3*, *ahk3-7*; *a4*, *cre1-2*; *a23*, *ahk2-5 ahk3-7*; *a24*, *ahk2-5 cre1-2*; *a34*, *ahk3-7 cre1-2*; *g14*, *abcg14*; *DM*, *cyp735a1-2 cyp735a2-2*; *a3g14*, *ahk3-7 abcg14*; *a3DM*, *ahk3-7 cyp735a1-2 cyp735a2-2*; *TCS*, *TCSn:GFP*; ns, not significant.

To determine which shoot-expressed CK receptor(s) mediate CK signaling underlying this growth effect, we analyzed grafted plants carrying single or double mutations in *AHK2*, *AHK3*, or *AHK4* specifically in WT/*ahk3* shoots. Grafted plants (scion/rootstock: WT/WT, WT/*ahk3*, *ahk2*/*ahk3*, *ahk3*/*ahk3*, *ahk4*/*ahk3*, *ahk23*/*ahk3*, *ahk24*/*ahk3*, *ahk34*/*ahk3*) were cultivated under 1/2 MS medium, and shoot growth was measured. Increases in shoot fresh weight and leaf blade area observed in WT/*ahk3* relative to WT/WT were similarly evident in *ahk2*/*ahk3*, *ahk4*/*ahk3*, and *ahk24*/*ahk3* (Fig. 6, C and D). In contrast, these enhancements were suppressed in *ahk3*/*ahk3* and *ahk34*/*ahk3*, and markedly reduced in *ahk23*/*ahk3*, where values were significantly lower than WT/WT (Fig. 6, C and D). These results indicate that enhanced shoot growth in WT/*ahk3* depends primarily on CK signaling through shoot-expressed AHK3, with partial contribution from AHK2. Next, to visualize sites of enhanced CK response in WT/*ahk3* shoots, we introduced the synthetic CK reporter *TCSn:GFP* into WT/WT and WT/*ahk3* shoots (Zürcher et al., 2013). Grafted plants (scion/rootstock: *TCSn:GFP*/WT, *TCSn:GFP*/*ahk3*) were grown on 1/2 MS medium. *TCSn:GFP*/*ahk3* exhibited greater shoot fresh weight and leaf blade area than *TCSn:GFP*/WT (Fig. 6E). Notably, GFP fluorescence intensity in true leaf was also significantly higher in *TCSn:GFP*/*ahk3* than in *TCSn:GFP*/WT (Fig. 6F). Finally, to examine how root-specific *AHK3* deficiency influences leaf micro-morphology, we measured epidermal cell size and number in true leaves of WT/WT and WT/*ahk3*. Cell size was comparable between genotypes, but cell number was significantly higher in WT/*ahk3* than in WT/WT (Fig. 6G). These results suggest that root-specific *AHK3* deficiency enhances leaf CK response, promoting epidermal cell division and thereby increasing leaf size.

### Root-expressed AHK3 modulates leaf size in a nitrate-dependent manner

*AHK3* was identified as one of the top genes co-expressed with the nitrate sensor NLP7 (Supplementary Table S1), suggesting that AHK3 may play a role in plant nitrate responses. To test this, we manipulated the nitrate levels of WT/WT and WT/*ahk3* roots. Ten-day-old WT/WT and WT/*ahk3* were first transferred to 2.5 mM ammonium succinate (5 mM NH_4_^+^) medium and incubated for 48 hours to deplete nitrate (Supplementary Fig. S11A). Grafted plants were then transferred to medium containing 5 mM NH_4_^+^ supplemented with either 0.5 mM KNO_3_ (KNO_3_) or 0.5 mM KCl (KCl) and grown for 5 days to assess the presence or absence of nitrate signaling in roots. Nitrate accumulated only in WT/WT and WT/*ahk3* roots grown under KNO_3_ (Supplementary Fig. S11B). Moreover, WT/WT and WT/*ahk3* appeared healthy, with comparable shoot chlorophyll concentrations under both conditions (Supplementary Fig. S12), indicating that nitrogen deficiency or ammonium toxicity did not occur. In WT/WT, shoot fresh weight and leaf blade area were found to be significantly greater under KNO_3_ than KCl (Fig. 7A), showing that nitrate signaling promotes leaf growth. Notably, shoot fresh weight and leaf blade area were similar between WT/WT and WT/*ahk3* under KCl, but were higher in WT/*ahk3* than in WT/WT under KNO_3_ (Fig. 7A).

**Figure 7.**
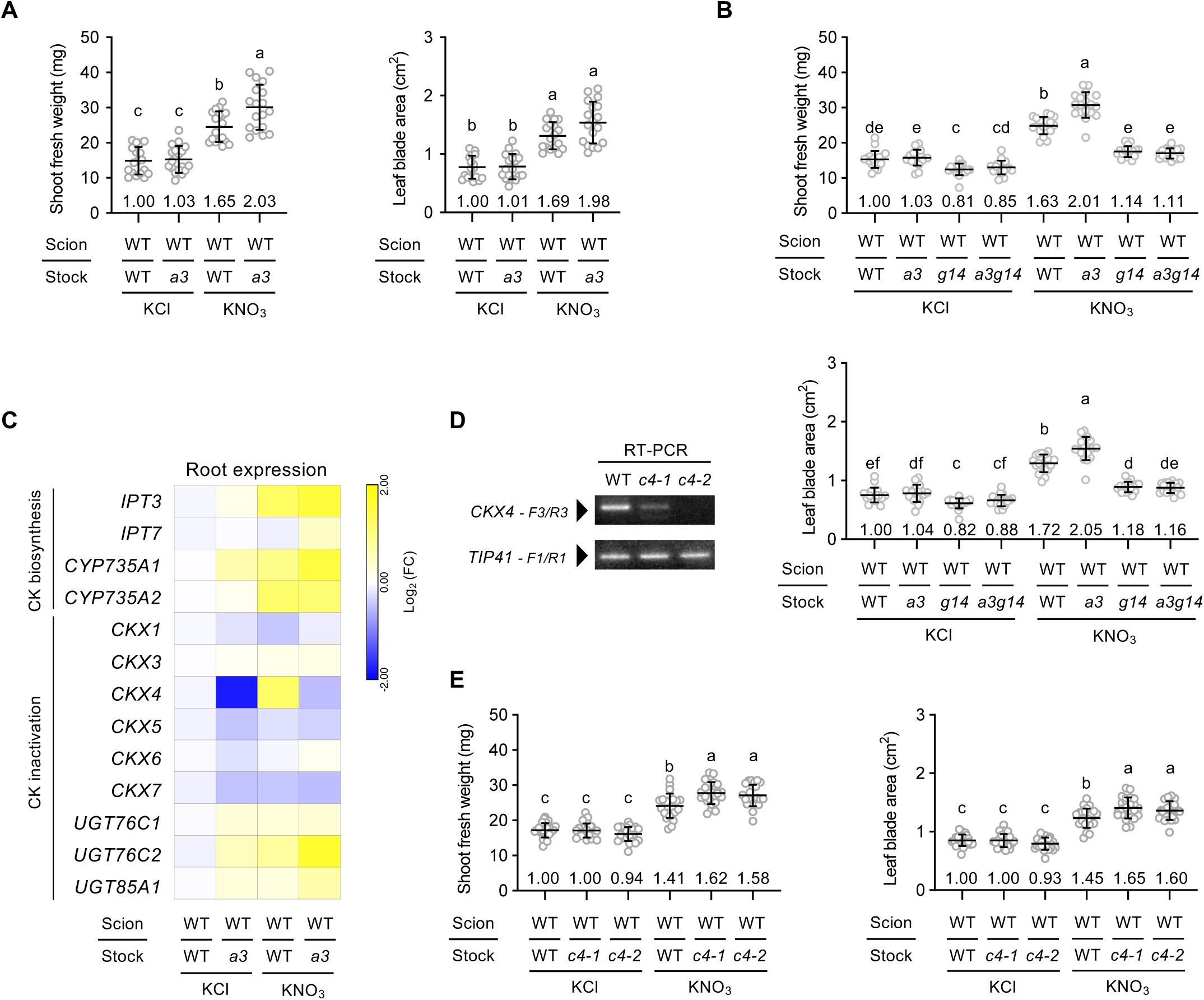
Effects of root-specific *AHK3* or *CKX4* deficiency on shoot growth in response to nitrate signaling. (A, B, E) Shoot fresh weight and leaf blade area of grafted plants 5 days after transfer from 5 mM NH_4_^+^ medium to either 5 mM NH_4_^+^ + 0.5 mM KNO_3_ or 5 mM NH_4_^+^ + 0.5 mM KCl: (A) WT/WT and WT/*ahk3*; (B) WT/WT, WT/*ahk3*, WT/*abcg14*, and WT/*ahk3 abcg14*; (E) WT/WT, WT/*ckx4-1*, and WT/*ckx4-2*. (C) Root expression of *IPT3*, *IPT7*, *CYP735A1*, *CYP735A2*, *CKX1*, *CKX3*, *CKX4*, *CKX5*, *CKX6*, *CKX7*, *UGT76C1*, *UGT76C2*, and *UGT85A1* 4 hours after transfer. Heatmaps were generated with MORPHEUS (https://software.broadinstitute.org/morpheus) from log_2_ fold change values relative to WT/WT under 5 mM NH_4_^+^ + 0.5 mM KCl (set = 1). Color scale: blue (minimum) to yellow (maximum) per row. (D) RT-PCR analysis of *CKX4* in WT, *ckx4-1*, and *ckx4-2* seedlings grown 7 days on 1/2 MS medium. *TIP41* was the control. Primers are listed in Supplementary Table S8. Data are shown as mean ± SD with replicates as follows: (A) n = 18, two replicates; (B) n = 18, three replicates; (C) n = 6, two replicates; (E) n = 24, four replicates. Different lowercase letters indicate significant differences, as determined via Tukey–Kramer test (*P* < 0.05). Graph numbers indicate mean values relative to WT/WT under 5 mM NH_4_^+^ + 0.5 mM KCl (set = 1). Grafted plants are denoted as “scion line/rootstock line.” WT, Col-0; *a3*, *ahk3-7*; *g14*, *abcg14*; *a3g14*, *ahk3-7 abcg14*; *c4-1*, *ckx4-1*; *c4-2*, *ckx4-2*; ns, not significant; KCl, 5 mM NH_4_^+^ + 0.5 mM KCl; KNO_3_, 5 mM NH_4_^+^ + 0.5 mM KNO_3_.

Next, to test whether nitrate-dependent growth promotion in WT/*ahk3* requires root-derived *t*Z-type CK signals, we analyzed WT/*abcg14* and WT/*ahk3 abcg14* grown under KNO_3_ or KCl. Interestingly, the increases in shoot fresh weight and leaf blade area observed in WT/WT and WT/*ahk3* under KNO_3_ were strongly suppressed in WT/*abcg14* and WT/*ahk3 abcg14* (Fig. 7B). These results indicate that long-distance signals of root-derived *t*Z-type CKs are required for nitrate-induced leaf growth promotion. Moreover, shoot fresh weight and leaf blade area were comparable between WT/*abcg14* and WT/*ahk3 abcg14* under KNO_3_ (Fig. 7B), suggesting that root-derived *t*Z-type CK signals may contribute to the enhanced leaf growth of WT/*ahk3*.

To further clarify the role of AHK3 in nitrate signaling, we analyzed expression of CK biosynthesis and inactivation genes in roots 4 hours after transfer to KNO_3_ or KCl. Nitrate accumulation was detected only in WT/WT and WT/*ahk3* roots after KNO_3_ treatment (Supplementary Fig. S11C). In WT/WT, root expression of *IPT3*, *CYP735A1*, *CYP735A2*, and *CKX4* was significantly higher under KNO_3_ than KCl (Fig. 7C, Supplementary Fig. S13), indicating induction by nitrate signaling. Importantly, expression of *IPT3*, *CYP735A1*, and *CYP735A2* was similar in WT/WT and WT/*ahk3* under KNO_3_, whereas *CKX4* expression was significantly reduced in WT/*ahk3* compared with WT/WT (Fig. 7C, Supplementary Fig. S13). These results suggest that both nitrate signals and root-expressed AHK3 are required for maximal *CKX4* induction.

To investigate the role of CKX4 in nitrate signaling, we obtained two independent T-DNA alleles, *ckx4-1* (SALK_055204) and *ckx4-2* (SALK_132516), from the Arabidopsis Biological Resource Center. *Ckx4-1* carries an intron insertion (Bartrina et al., 2011), while *ckx4-2* carries an exon insertion (Supplementary Fig. S14). RT-PCR confirmed reduced *CKX4* transcript in *ckx4-1* and absence in *ckx4-2* (Fig. 7D), indicating that *ckx4-1* is a knockdown and *ckx4-2* a knockout. Grafted plants (scion/rootstock: WT/WT, WT/*ckx4-1*, WT/*ckx4-2*) were cultivated under KNO_3_ or KCl, and shoot growth was measured. Under KCl, shoot fresh weight and leaf blade area were similar among genotypes (Fig. 7E). Under KNO_3_, however, both traits were significantly greater in WT/*ckx4-1* and WT/*ckx4-2* compared to WT/WT (Fig. 7E).

To further clarify relationship between AHK3 and CKX4, we generated *ahk3 ckx4-1* and *ahk3 ckx4-2* mutants by traditional crossing (Supplementary Fig. S15, A and B). Grafted plants (scion/rootstock: WT/WT, WT/*ahk3*, WT/*ckx4-1*, WT/*ckx4-2*, WT/*ahk3 ckx4-1*, WT/*ahk3 ckx4-2*) were grown under KNO_3_ or KCl, and shoot growth was measured. All genotypes showed comparable shoot growth under KCl (Supplementary Fig. S15C). Under KNO_3_, however, shoot fresh weight and leaf blade area were significantly greater in WT/*ahk3*, WT/*ckx4-1*, WT/*ckx4-2*, WT/*ahk3 ckx4-1*, and WT/*ahk3 ckx4-2* compared with WT/WT (Supplementary Fig. S15C). These traits were significantly higher in WT/*ahk3* than in WT/*ckx4-1* and WT/*ckx4-2* (Supplementary Fig. S15C), indicating that CKX4 acts downstream of AHK3. Importantly, WT/*ahk3 ckx4-1* and WT/*ahk3 ckx4-2* showed no further growth increase relative to WT/*ahk3* under KNO_3_ (Supplementary Fig. S15C). Together, these results suggest that nitrate-induced leaf growth promotion occurs through a pathway mediated jointly by AHK3 and CKX4.

## DISCUSSION

Recent studies have shown that long-distance signals of root-derived *t*Z-type CK coordinate multiple physiological processes in aerial organs, including leaf growth, photoperiod stress response, shade avoidance, and flowering time (Osugi et al., 2017; Landrein et al., 2018; Frank et al., 2020; Gautrat et al., 2024; Bartrina et al., 2025). These findings highlight that proper modulation of root-to-shoot *t*Z-type CK transport is essential for plants to adapt to environmental fluctuations.

Based on our findings and prior studies (Miyawaki et al., 2004; Kiba et al., 2013; Ko et al., 2014; Zhang et al., 2014; Sakakibara, 2021, 2025), we provide new insight into the specific mechanisms responsible for modulating xylem *t*Z-type CK transport, as mediated by CK perception and degradation through an AHK3–CKX4 module. Nitrate signaling promotes *t*Z-type CK biosynthesis in pericycle and phloem companion cells via IPT3, CYP735A2, and related enzymes. *t*Z-type CKs are then transported to the extracellular and endoplasmic reticulum (ER) space by transporters such as ABCG14. Active form *t*Z are perceived by AHK3 on plasma membrane (PM) and ER membrane. Activation of CK signaling through AHK3 induces expression of *CKX4*, predicted to localize in apoplast, where it reduces apoplastic *t*Z-type CK levels by degradation. These mechanisms likely influence the apoplastic *t*Z-type CK pool and, consequently, the amount of *t*Z-type CK loaded into xylem (Supplementary Fig. S16). To further substantiate this model, it will likely be necessary to analyze the spatial expression patterns of AHK3 and CKX4 at the tissue and cellular level in future studies. Interestingly, *IPT3* expression and *t*Z-type CK levels in roots also respond to other nutrients, including phosphate, sulfate, and potassium (Hirose et al., 2008; Nam et al., 2012). Thus, AHK3 may modulates *t*Z-type CK levels in response to diverse nutrient signals. Collectively, the AHK3–CKX4 module could serve as a feedback regulator that attenuates the nutrient-induced xylem transport of *t*Z-type CKs, thereby fine-tuning leaf CK response to maintain balanced plant growth.

Root-specific deficiency of *AHK3* enhanced shoot growth but decreased primary root growth, resulting in an imbalance in whole-plant growth (Fig. 1, Supplementary Fig. S2). Given that primary root length in *ahk3* is comparable to that in Col-0 (Riefler et al., 2006), the reduced root growth observed in WT/*ahk3* is considered a compensatory response to enhanced shoot growth. Interestingly, a temporal analysis of shoot growth in grafted plants revealed that WT/*ahk3* exhibited a significant increase in leaf blade area 3 days after transfer to 1/2 MS medium, which coincided with the onset of primary root growth inhibition (Supplementary Figs. S2 and S17). Furthermore, not only was shoot growth enhancement attenuated in *ahk3*/*ahk3*, but primary root growth inhibition was also diminished (Fig. 6, Supplementary Fig. S17), suggesting that shoot and root growth may change in a correlated manner. Taken together, these results indicate that root-expressed AHK3 is physiologically important for maintaining whole-plant growth balance by modulating xylem *t*Z-type CK transport and leaf CK status, thereby preventing excessive energy investment in shoot growth and coordinating energy allocation between shoots and roots.

CKXs are known to mediate irreversible CK degradation, producing adenine (or its derivatives) and aldehyde (Schmülling et al., 2003). In *Arabidopsis*, seven CKXs differ in substrate specificity, spatial and temporal expression, and subcellular localization, targeting either vacuoles (CKX1, CKX3), ER (CKX1), apoplasts (CKX2, CKX4, CKX5, CKX6), or the cytosol (CKX7) (Werner et al., 2003; Galuszka et al., 2007; Frébort et al., 2011; Niemann et al., 2018). Expression of *CKX3*, *CKX4*, *CKX5*, and *CKX6* is strongly induced by CK treatment (Werner et al., 2006), suggesting sensitivity to internal CK levels and roles in maintaining homeostasis. Moreover, CKX2 and CKX4 show higher *in vitro* CKX activity than other CKXs, preferentially degrading active form CKs at neutral or slightly basic pH (Galuszka et al., 2007). Our data showed that root-specific *AHK3* deficiency greatly reduced *CKX4* expression in roots, without significantly altering other *CKX*s (Figs. 3 and 7). These findings suggest that root-expressed AHK3 modulates *t*Z degradation by inducing *CKX4* expression in response to the internal *t*Z level. This interpretation is supported by our observation that *t*Z concentration was markedly higher in WT/*ahk3* xylem sap than in WT/WT (Fig. 4). Furthermore, Osugi *et al*. (2017) demonstrated that root-derived *t*ZR controls both leaf number and area, whereas root-derived *t*Z regulates only leaf area. In WT/*ahk3*, the increase in xylem sap *t*Z relative to WT/WT was greater than that of *t*ZR (Fig. 4). Thus, the enlarged leaf blade area in WT/*ahk3*, without a change in leaf number, likely reflects long-distance signaling by root-derived *t*Z (Figs. 1 and 4).

AHK2, AHK3, and AHK4 localize to the PM and ER membrane, where they play central roles in CK signal transduction (Antoniadi et al., 2020; Kubiasová et al., 2020). AHK3 is considered to be primarily localized on the ER membrane and partially on the PM (Caesar et al., 2011; Wulfetange et al., 2011). Therefore, ER-localized AHK3 may contribute substantially to modulation of xylem transport of *t*Z-type CKs. AHK3 is also expressed more strongly in leaves than in roots, and its leaf expression exceeds that of AHK2 and AHK4, supporting its hypothesized role as a major regulator of leaf CK response (Higuchi et al., 2004). This hypothesis is consistent with our finding that leaf growth promotion, driven by increased shoot CK response, was diminished when AHK3 was specifically disrupted in WT/*ahk3* shoots (Fig. 6). Strikingly, root-expressed AHK3 exerted a stronger impact on the root transcriptome and on CK levels than root-expressed AHK2 and AHK4 (Figs. 2 and 3). Taken together, these results suggest that AHK3 also plays a major role in roots. Previous reports indicate that rosette leaf growth, CK metabolism, leaf senescence, stomatal opening, and embryo growth depend particularly on AHK3, as disruption of AHK3, but not other CK receptors, impairs these processes (Riefler et al., 2006; Kim et al., 2006; Arnaud et al., 2017; Loperfido et al., 2025). The distinct functions of these receptors may arise from their expression patterns, recognition of different CK species, and specific interactions with other proteins (Higuchi et al., 2004; Spíchal et al., 2004; Romanov et al., 2006; Stolz et al., 2011). A key feature of AHK3 is its ligand preference: while AHK2 and AHK4 bind iP and *t*Z with similar affinity, AHK3 shows higher affinity for *t*Z than iP (Stolz et al., 2011). In addition, AHK3 specifically mediates CK-triggered phosphorylation of ARR2 (Kim et al., 2006). These differences in ligand preference and downstream targets may underlie the unique functions of AHK3.

Overall, this study uncovered a comprehensive mechanism by which the CK receptor AHK3 systemically modulates the shoot growth via exerting control of xylem *t*Z-type CK transport. Furthermore, we clarified the role of AHK3 in modulating leaf size in response to root nitrogen availability, a key environmental factor. Overall, these findings provide new insight into how plants integrate nutritional cues to coordinate systemic communication between roots and shoots.

## MATERIALS AND METHODS

### Plant materials

The *Arabidopsis thaliana* accession Col-0 was used as wild type. T-DNA insertion lines *ahk2-5*, *ahk3-7*, *cre1-2*, *ahk2-5 ahk3-7*, *ahk2-5 cre1-2*, *ahk3-7 cre1-2* (Riefler et al., 2006), *abcg14* (Ko et al., 2014), *cyp735a1-2 cyp735a2-2* (Kiba et al., 2013), and *ckx4-1* (Bartrina et al., 2011) have been described previously. The *ckx4-2* (SALK_132516) line was obtained from the Arabidopsis Biological Resource Center (https://abrc.osu.edu/). The transgenic line *TCSn:GFP* (Zürcher et al., 2013) was described previously. Finally, the homozygous multiple mutants *ahk3-7 abcg14*, *ahk3-7 cyp735a1-2 cyp735a2-2*, *ahk3-7 ckx4-1*, and *ahk3-7 ckx4-2* were generated by crossing, and genotypes were determined by genomic PCR using primers listed in Supplementary Table S8.

### Growth conditions of grafted plants

Seeds were surface-sterilized and kept in the dark at 4°C for 3 days to break dormancy. Approximately 120 seeds per dish were sown in rectangular dishes (140 × 100 × 20 mm^3^; Eiken Chemical Co. Ltd., Tokyo, Japan) containing 50 mL of half-strength MS salts including vitamins (Code: M0222; Duchefa Biochemie), supplemented with 0.05% (w/v) MES-KOH (pH 5.7), 1% (w/v) sucrose, and 0.5% (w/v) gellan gum (Fujifilm Wako, VA, USA). Plants were grown under a 16/8-h light/dark cycle at 22°C with a photosynthetic photon flux density (PPFD) of 25 μmol m^-2^ s^-1^. Five-day-old seedlings were cut perpendicularly at the hypocotyl using an injection needle tip (Code: NN-2613S; TERUMO, Tokyo, Japan). Excised scions were connected to corresponding rootstocks using silicone microtubes (inner × outer diameter: φ0.4 × φ0.5 mm; Code: 1-8194-04; AS ONE, Osaka, Japan). Grafted plants were incubated for 5 days under continuous light at 27°C with a PPFD of 50 μmol m^-2^ s^-1^. Successfully grafted plants without adventitious roots were selected, transferred to solid media or soil pots, then grown for further experiments.

For *in vitro* cultures, grafted plants were grown vertically for five days or horizontally for seven days under a PPFD of 70 μmol m^-2^ s^-1^ (16/8-h light/dark cycle) at 22°C in 50 mL (vertical) or 30 mL (horizontal) of half-strength MS salts without vitamins (Code: M0221; Duchefa Biochemie), supplemented with 0.05% (w/v) MES-KOH (pH 5.7), 1% (w/v) sucrose, and 0.8% (w/v) purified agar (Code: 01162-15; Nacalai Tesque, Kyoto, Japan). To facilitate non-destructive root sampling during horizontal growth, media surfaces were covered with cellophane sheets (Hachiya et al., 2021).

For soil cultivation, grafted plants were grown for 7 days under a PPFD of 70 μmol m^-2^ s^-1^ (16/8-h light/dark cycle) at 22°C in pots containing ∼150 mL of nutrient-rich soil (Supermix A; Sakata Seed, Kyoto, Japan).

For transfer experiments with varying nitrogen concentrations, grafted plants were placed on cellophane sheets in round dishes (diameter 90 mm; depth 20 mm; AS ONE) containing 30 mL of half-strength MS modified basal salt mixture without nitrogen (M531; PhytoTech Labs, Lenexa, KS, USA), supplemented with 2.5 mM (NH_4_)_2_C_4_H_4_O_4_, 0.05% (w/v) MES-KOH (pH 5.7), 1% (w/v) sucrose, and 0.8% (w/v) purified agar (Code: 01162-15; Nacalai Tesque) (5 mM NH_4_^+^ medium). Plants were grown for 48 hours under a PPFD of 70 μmol m^-2^ s^-1^ (16/8-h light/dark cycle) at 22°C, then transferred to 5 mM NH_4_^+^ medium supplemented with either 0.5 mM KNO_3_ or 0.5 mM KCl and grown for five days under the same conditions.

### Growth analysis of grafted plants

Shoots were weighed on a precision balance (HR-202i; A&D, Tokyo, Japan). All leaf blades were harvested and scanned at 600 dpi (GT-X980; EPSON, Tokyo, Japan) to measure true leaf blade area. Roots of vertically grown plants were scanned at 600 dpi to measure primary root length. To monitor root elongation, root tip positions were marked every 24 hours. Leaf blade area and primary root length were quantified using ImageJ version 1.53t (US National Institutes of Health, Bethesda, MD, USA).

### Xylem sap collection

Grafted plants were cut perpendicularly at the hypocotyl using an injection needle tip (Code: NN-2613S; TERUMO). A silicone microtube (inner × outer diameter: φ0.5 × φ0.6 mm; Code: 1-8194-05; AS ONE) was attached to the cut end, and plants were incubated for 2 hours to allow exudation. After incubation, the silicone microtube was detached, and xylem sap collected in the tube was recovered by centrifugation into a collection tube. This was then and stored at –80°C until further analysis. We used 12 µL aliquots of sap from each grafted plant for CK analyses.

### Determination of CK species

CK extraction and quantification followed the method of Kojima *et al*. (2009), using an ultra-performance liquid chromatography–tandem quadrupole mass spectrometry system (AQUITY UPLC System/XEVO-TQS; Waters Corp., Milford, MA, USA) equipped with an octadecylsilyl column (ACQUITY UPLC HSS T3, 1.8 µm, 2.1 mm × 100 mm; Waters Corp.).

### Extraction of total RNA

To extract total RNA, frozen plant samples were first homogenized using a Multi-Beads Shocker and 5 mm zirconia beads. Total RNA was then isolated with the RNeasy Plant Mini Kit (Qiagen, Hilden, Germany) with all protocols performed as per the manufacturer’s protocol. For RNA-seq library preparation, RNA was further purified by on-column DNase digestion (Qiagen).

### RNA-seq analysis

RNA quality was evaluated using a Qubit RNA IQ Assay Kit with a Qubit 4 Fluorometer (Thermo Fisher Scientific, Tokyo, Japan). Samples with RNA IQ scores of 9.5–10.0 were used for RNA-seq. cDNA libraries were prepared with the NEBNext Ultra II Directional RNA Library Prep Kit (E7760S, New England Biolabs, Ipswich, MA, USA), SPRIselect (B23317, Beckman Coulter), the NEBNext Poly(A) mRNA Magnetic Isolation Module (E7490S, New England Biolabs), and NEBNext Multiplex Oligos for Illumina (E7710S, New England Biolabs), with all steps following the manufacturer’s instructions. Libraries were sequenced on a NextSeq 500 (Illumina, San Diego, CA, USA), and bcl files were converted to fastq using bcl2fastq (Illumina). Reads were analyzed as described by Notaguchi *et al*. (2014) and mapped to the *Arabidopsis* reference (TAIR10) using Bowtie (Langmead et al., 2009) with the options “--all --best --strata.” For normalization, read count data were transformed with rlog and adjusted by subtracting the average expression for each gene using default settings of iDEP v0.96 (Ge et al., 2018). DEGs were extracted using edgeR with a false discovery rate cutoff of 0.15 under the default settings of RNAseqChef v1.1.4 (Etoh et al., 2023). Gene set enrichment was performed using ShinyGO v0.82 (Ge et al., 2020) using the default parameters. During analysis we focused on the Biological Process annotation category.

### Reverse transcription and quantitative RT-PCR

Reverse transcription was performed with the ReverTraAce qPCR RT Master Mix with gDNA Remover (Toyobo, Osaka, Japan). For each reaction, 0.5 µg of RNA was used as input. Synthesized cDNA was diluted five-fold with sterile water and used as template for qRT-PCR. Reactions were run on a QuantStudio 1 system (Thermo Fisher Scientific) using a KOD SYBR qPCR Mix (Toyobo). Relative expression was determined using the comparative Ct method. *TIP41* was used as an internal reference gene. Primer sequences are listed in Supplementary Table S9.

### Determination of nutrient elements

Shoots were harvested and oven-dried at 80°C for 2 days, ground with a spoon, then stored in a desiccator with silica gel. For carbon and nitrogen quantification, we weighed portions of dried samples, packed them into tin capsules, and subjected them to quantitative analysis using a vario MICRO cube elemental analyzer (Elementar Analysensysteme GmbH, Hanau, Germany). For other elements (e.g., potassium, calcium, sulfur, phosphorus, magnesium, sodium, manganese, zinc, iron, boron, copper, molybdenum, and cobalt), weighed portions of dried samples were digested with HNO_3_ followed by H_2_O_2_. Digests were suspended in 1 mL of 0.08 M HNO_3_, diluted tenfold with 0.08 M HNO_3_, and analyzed using inductively coupled plasma mass spectrometry (Agilent 7800 ICP-MS; Agilent Technologies, Santa Clara, CA, USA). Here, indium was used as an internal standard.

### Nitrate level determination

Nitrate was extracted using 10 volumes of sterile water at 100°C for 20 min. A 5.0 µL aliquot of supernatant was mixed with 40 µL of 5% (w/v) salicylic acid in concentrated sulfuric acid and incubated at room temperature for 20 min. A mock treatment with 40 µL concentrated sulfuric acid was prepared in parallel. Finally, 1 mL of 8% (w/v) NaOH was added to the mixture, and nitrate concentrations was determined by recording sample absorbance at 410 nm.

### Fluorescence microscopy

Fluorescence imaging was performed using an all-in-one fluorescence microscope (BZ-X710, KEYENCE, Osaka, Japan) equipped with a 4× objective (CFI Plan Fluor DL 4×/0.13, Nikon, Tokyo, Japan) and a metal halide lamp (No. 91056, KEYENCE). A narrow-band GFP filter (Ex. 480/20; Em. 510/20; 49020, Chroma, Brattleboro, VT, USA) was used for GFP detection, and a Cy5 filter (Ex. 620/60; Em. 700/75; 49006, Chroma) was used for chlorophyll autofluorescence. All images were acquired using identical exposure times. For quantification of GFP fluorescence, background signals were subtracted by using non-fluorescent regions as reference.

### Leaf cellular analysis

The fifth true leaf was excised from the shoot and scanned at 600 dpi (GT-X980; EPSON) to measure total leaf area. The abaxial surface was pressed onto Scotch transparent adhesive tape (Code: 500-3-15-10P; 3M Japan, Tokyo, Japan) and carefully peeled to isolate epidermal cells. Epidermal cell images were captured using a fluorescence microscope (BZ-X710; KEYENCE) equipped with a 20× objective (CFI Plan Apo 20×/0.75; Nikon). Thirty epidermal cells were randomly selected, and their areas were measured using ImageJ v1.53t (US National Institutes of Health, Bethesda). The mean area of 30 cells was defined as epidermal cell size. The total epidermal cell number was estimated by dividing the total leaf area by the mean cell size.

### Statistical analyses

Data were analyzed using unpaired two-tailed Welch’s *t*-tests and Tukey–Kramer multiple comparison test. All statistical analyses were conducted in R v4.3.0 (R Foundation for Statistical Computing, Vienna, Austria).

## AUTHOR CONTRIBUTIONS

K.M. and T.H. conceived and designed experiments. K.M., T.S., M.K., Y.T., T.Ka., H.S., and T.H. performed the experiments and data analysis. T.Ki., T.N., and T.H. supervised the study. K.M. wrote the manuscript with contributions from all authors. All authors approved the final manuscript.

## ACKNOWLEDGEMENTS

We thank Dr. Ryo Tabata and Dr. Tomoko Niwa for technical advice, Ms. Maki Saiki for ICP-MS analysis, and Prof. Thomas Schmülling for important input. We also acknowledge the use of ChatGPT, an AI-based language tool, for support in clarifying the manuscript, and Enago (www.enago.jp) for English language review. This study was supported by grants from JSPS KAKENHI [No. JP24KJ1709 to K.M., No. 21H02087 to T.Ka., and Nos. 20K05771 and 23K04978 to T.H.] and from JST SPRING [No. JPMJSP2155 to K.M.].

## DATA AVAILABILITY STATEMENT

RNA-seq raw data are deposited in ArrayExpress under accession number E-MTAB-15416.

## SUPPORTING INFORMATION

Additional Supporting Information is found in the online version of this article.

**Figure S1.** Effects of root-specific CK receptor deficiency on shoot growth in soil.

**Figure S2.** Effects of root-specific CK receptor deficiency on primary root length.

**Figure S3.** Effects of root-specific CK receptor deficiency on root concentration of individual CK species.

**Figure S4.** Differentially expressed genes in grafted plants visualized by volcano plot.

**Figure S5.** Differentially expressed genes in grafted plants visualized by Venn diagram.

**Figure S6.** Differentially expressed genes in grafted plants visualized by gene ontology terms.

**Figure S7.** Effects of root-specific CK receptor deficiency on xylem sap concentration of individual CK species.

**Figure S8.** Effects of root-specific CK receptor deficiency on shoot concentration of individual CK species.

**Figure S9.** Effects of root-specific CK receptor deficiency on shoot element concentrations.

**Figure S10.** Genomic PCR and RT-PCR results for *ahk3 abcg14* and *ahk3 cypDM*.

**Figure S11.** Effects of root-specific *AHK3* deficiency on root nitrate concentration in response to nitrate signaling.

**Figure S12.** Effects of root-specific *AHK3* deficiency on shoot chlorophyll concentration in response to nitrate signaling.

**Figure S13.** Effects of root-specific *AHK3* deficiency on root gene expressions in response to nitrate signaling.

**Figure S14.** Characterization of *CKX4* T-DNA insertion alleles.

**Figure S15.** Effects of root-specific deficiencies of *AHK3* and *CKX4* on shoot growth in response to nitrate signaling.

**Figure S16.** A proposed model for xylem *t*Z-type CK transport controlled by AHK3 in response to nitrate signaling.

**Figure S17.** Effects of root-specific *AHK3* deficiency on shoot and root growth.

**Table S1.** A list coexpressed with *NLP7* by ATTED II.

**Table S2.** Information about raw read counts.

**Table S3.** Information about reads per million mapped reads.

**Table S4.** Information about root normalized transcript levels.

**Table S5.** Information about shoot normalized transcript levels.

**Table S6.** Information about root DEGs.

**Table S7.** Information about shoot DEGs.

**Table S8.** Primers used in this study for genomic PCR and RT-PCR.

**Table S9.** Primers used in this study for qRT-PCR.

