## Supplemental Data 1 for "Cytokinin receptor AHK3 influences leaf size by modulating *trans*-zeatin-type cytokinin levels in xylem"

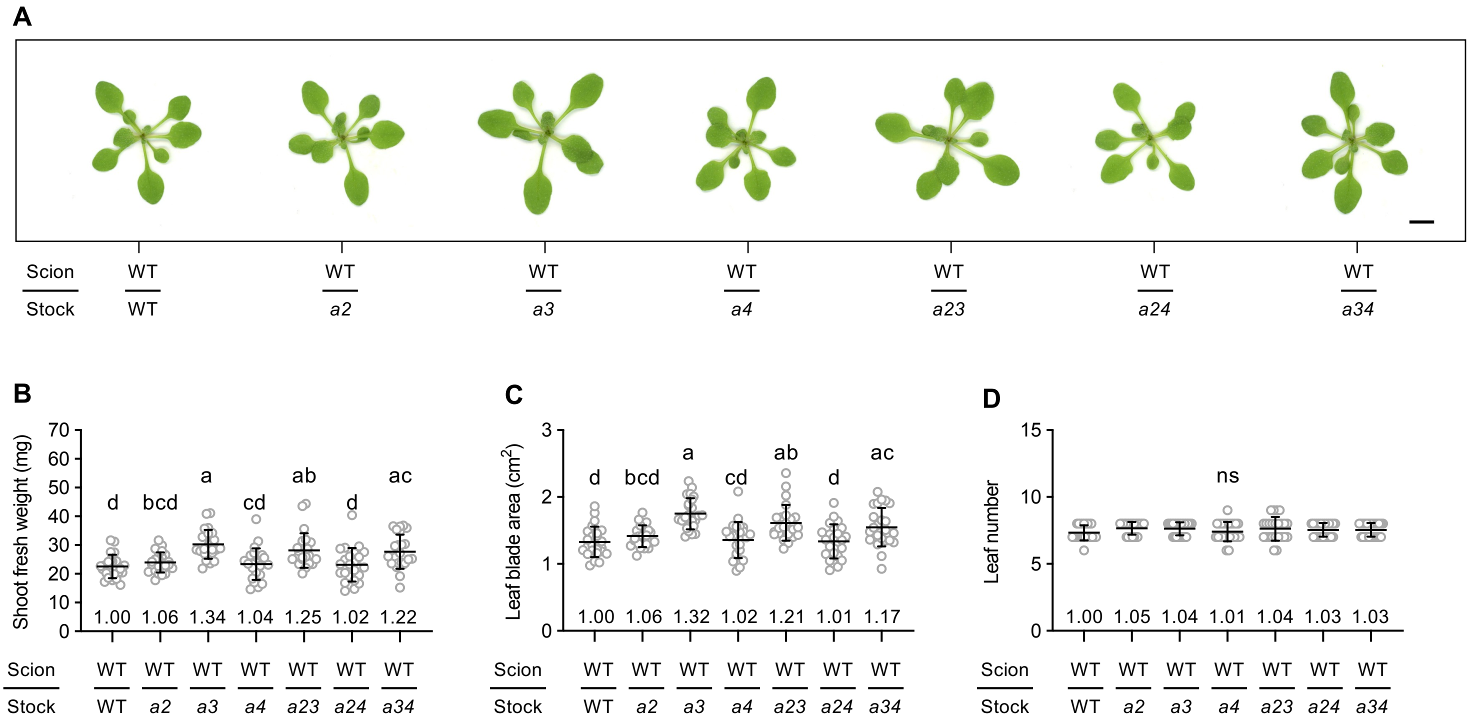


**Figure S1. Effects of root-specific CK receptor deficiency on shoot growth in soil.** (A) Representative shoots of grafted plants. Scale bars = 5 mm. (B–D) Shoot fresh weight (B), leaf blade area (C), and leaf number (D) of grafted plants. Data pooled from six independent grafting experiments are presented as mean ± SD (n = 24). Different lowercase letters indicate significant differences, as determined via Tukey–Kramer test ($P < 0.05$). Numbers on graphs show mean values relative to WT/WT (set = 1). Grafted plants are denoted as "scion line/rootstock line." WT, Col-0; *a2*, *ahk2-5*; *a3*, *ahk3-7*; *a4*, *cre1-2*; *a23*, *ahk2-5 ahk3-7*; *a24*, *ahk2-5 cre1-2*; *a34*, *ahk3-7 cre1-2*; ns, not significant.

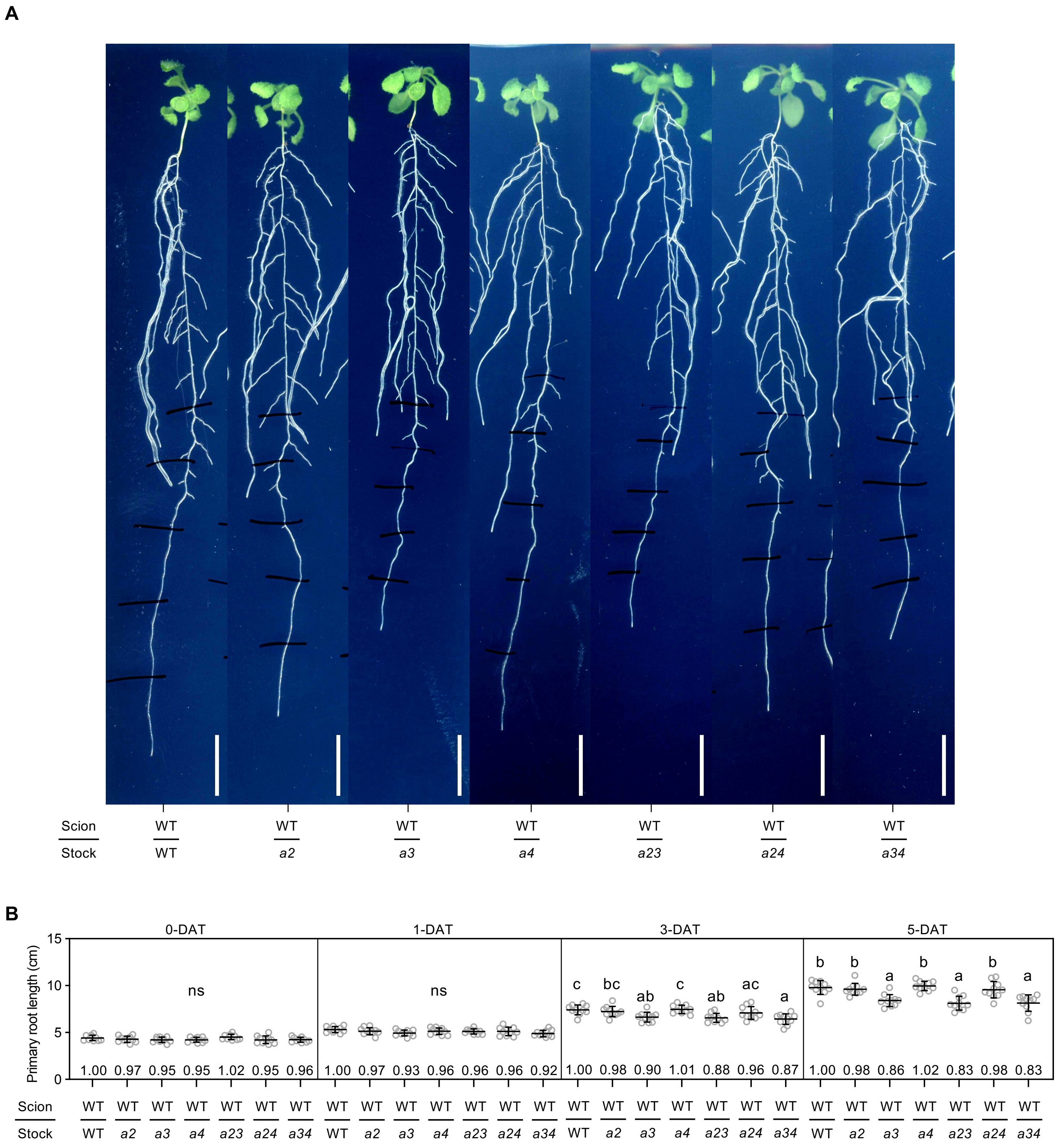


**Figure S2. Effects of root-specific CK receptor deficiency on primary root length.** (A) Representative roots of grafted plants 5 days after transfer 1/2 MS medium. Scale bars = 1 cm. (B) Time course of primary root length in grafted plants after transfer to 1/2 MS medium. Data are means ± SD (n = 10). Different lowercase letters indicate significant differences, as determined via Tukey–Kramer test ($P < 0.05$). Numbers on graphs show mean values relative to WT/WT (set = 1). Grafted plants are denoted as "scion line/rootstock line." WT, Col-0; *a2*, *ahk2-5*; *a3*, *ahk3-7*; *a4*, *cre1-2*; *a23*, *ahk2-5 ahk3-7*; *a24*, *ahk2-5 cre1-2*; *a34*, *ahk3-7 cre1-2*; ns, not significant; DAT, days after transfer.

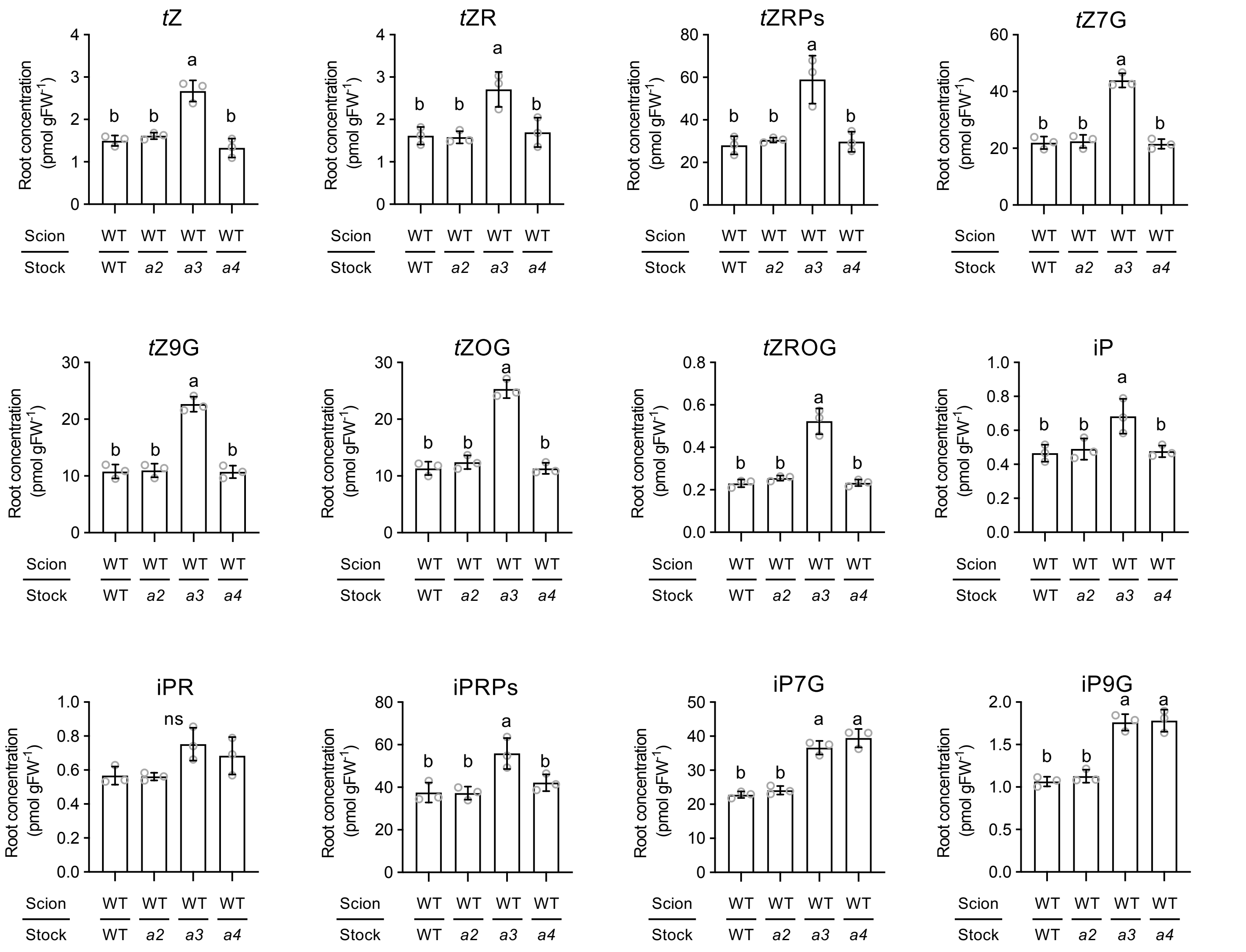


**Figure S3. Effects of root-specific CK receptor deficiency on root concentration of individual CK species.** Root concentrations of *t*Z, *t*ZR, *t*ZRPs, *t*Z7G, *t*Z9G, *t*ZOG, *t*ZROG, iP, iPR, iPRPs, iP7G, and iP9G in grafted plants. Data pooled from three independent grafting experiments are presented as mean ± SD (n = 3). Different lowercase letters indicate significant differences, as determined via Tukey–Kramer test ($P < 0.05$). Grafted plants are denoted as “scion line/rootstock line.” WT, Col-0; *a2*, *ahk2-5*; *a3*, *ahk3-7*; *a4*, *cre1-2;* FW, fresh weight; *t*Z, *trans*-zeatin; *t*ZR, *t*Z-riboside; *t*ZRPs, *t*ZR 5′-phosphates; *t*Z7G, *t*Z-7 *N*-glucoside; *t*Z9G, *t*Z-9 *N*-glucoside; *t*ZOG, *t*Z-*O*-glucoside; *t*ZROG, *t*ZR-*O*-glucoside; iP, $N^6$-($\Delta^2$-isopentenyl) adenine; iPR, iP-riboside; iPRPs, iPR 5′-phosphates; iP7G, iP-7 *N*-glucoside; iP9G, iP-9 *N*-glucoside; ns, not significant.

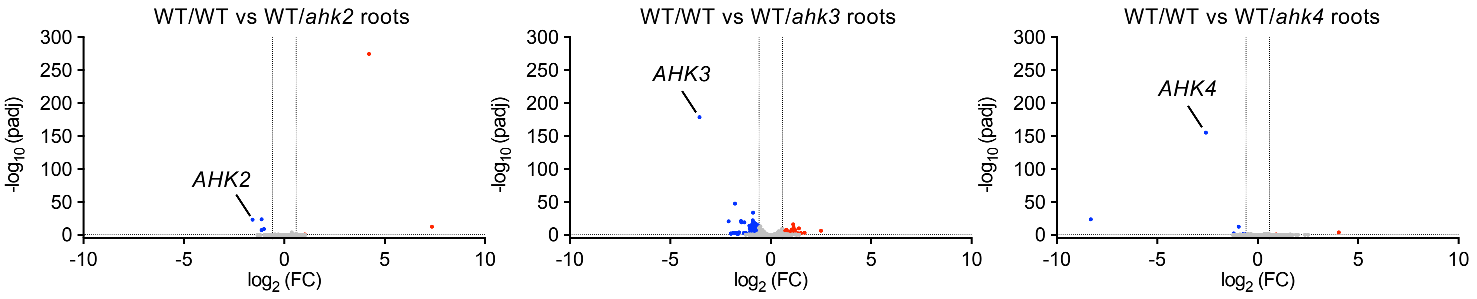


**Figure S4. Differentially expressed genes in grafted plants visualized by volcano plot.** Volcano plots showing the full range of DEGs between WT/WT and WT/*ahk2*, WT/*ahk3*, or WT/*ahk4* roots. DEG analysis was performed with RNAseqChef (Etoh et al., 2023) using edgeR, applying a 1.5-fold change threshold and FDR cutoff of 0.15 (all other parameters default). x-axis: $\log_2$ fold change ($\log_2$ FC); y-axis: $-\log_{10}$ adjusted p-value (padj). Each dot represents a single gene. Red, blue, and grey indicate upregulated, downregulated, and non-significant genes, respectively. Grafted plants are denoted as “scion line/rootstock line.” WT, Col-0; FC, fold change.

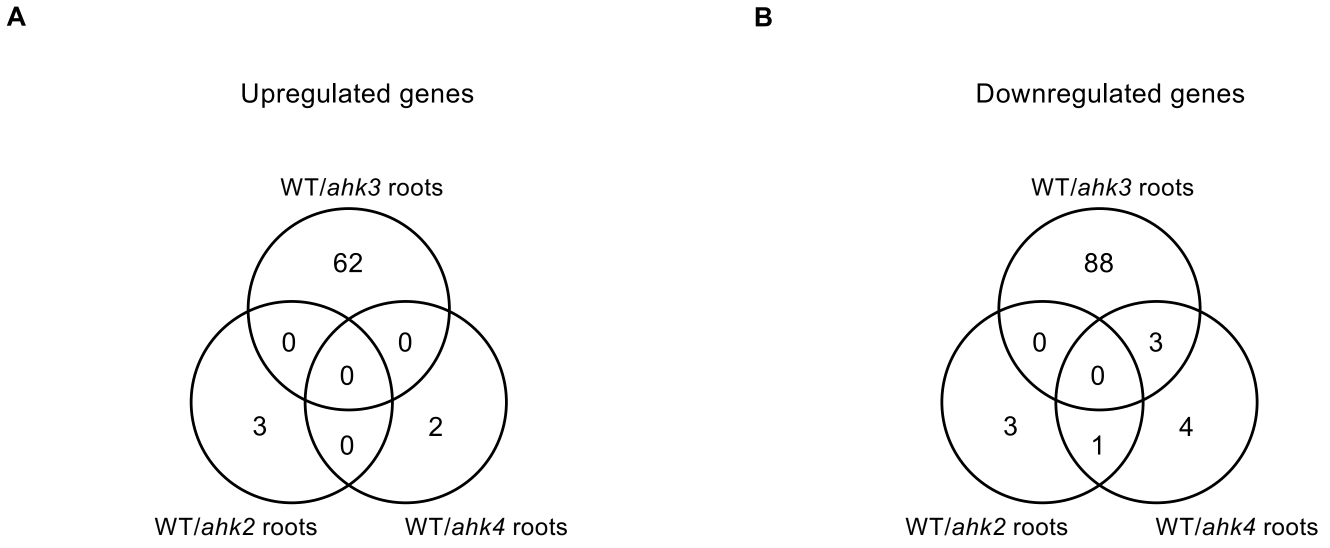


**Figure S5. Differentially expressed genes in grafted plants visualized by Venn diagram.** (A–B) Venn diagrams showing the overlap of upregulated (A) and downregulated (B) genes between WT/WT and WT/*ahk2*, WT/*ahk3*, or WT/*ahk4* roots. DEG analysis was performed with RNAseqChef (Etoh et al., 2023) using edgeR, applying a 1.5-fold change threshold and FDR cutoff of 0.15 (all other parameters default).

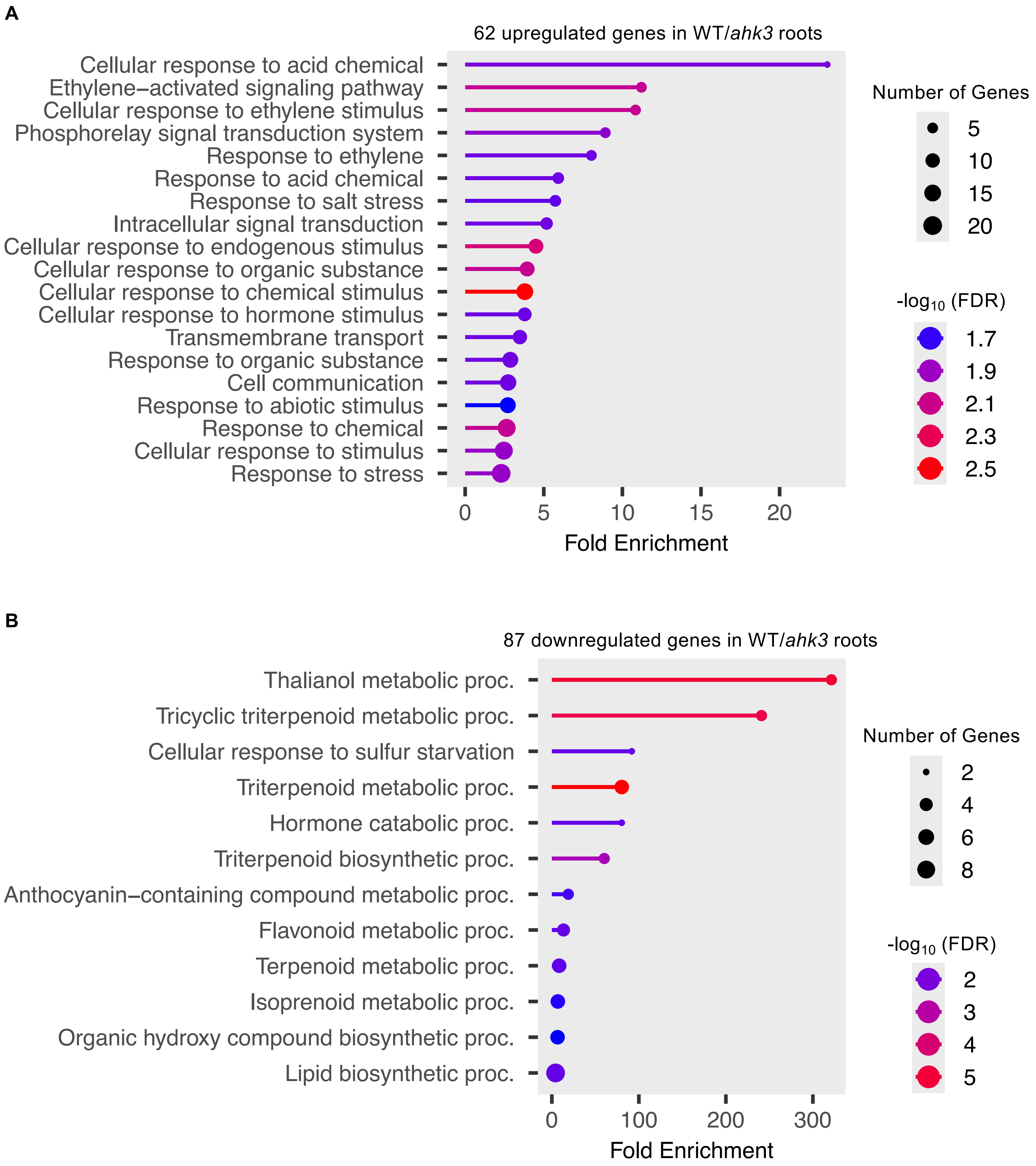


**Figure S6. Differentially expressed genes in grafted plants visualized by gene ontology terms.** (A–B) Lollipop plots showing GO enrichment analysis from the 62 genes upregulated (A) and 87 genes downregulated (B) in WT/*ahk3* roots, generated with ShinyGO (Ge et al., 2020) using default settings for biological process terms.

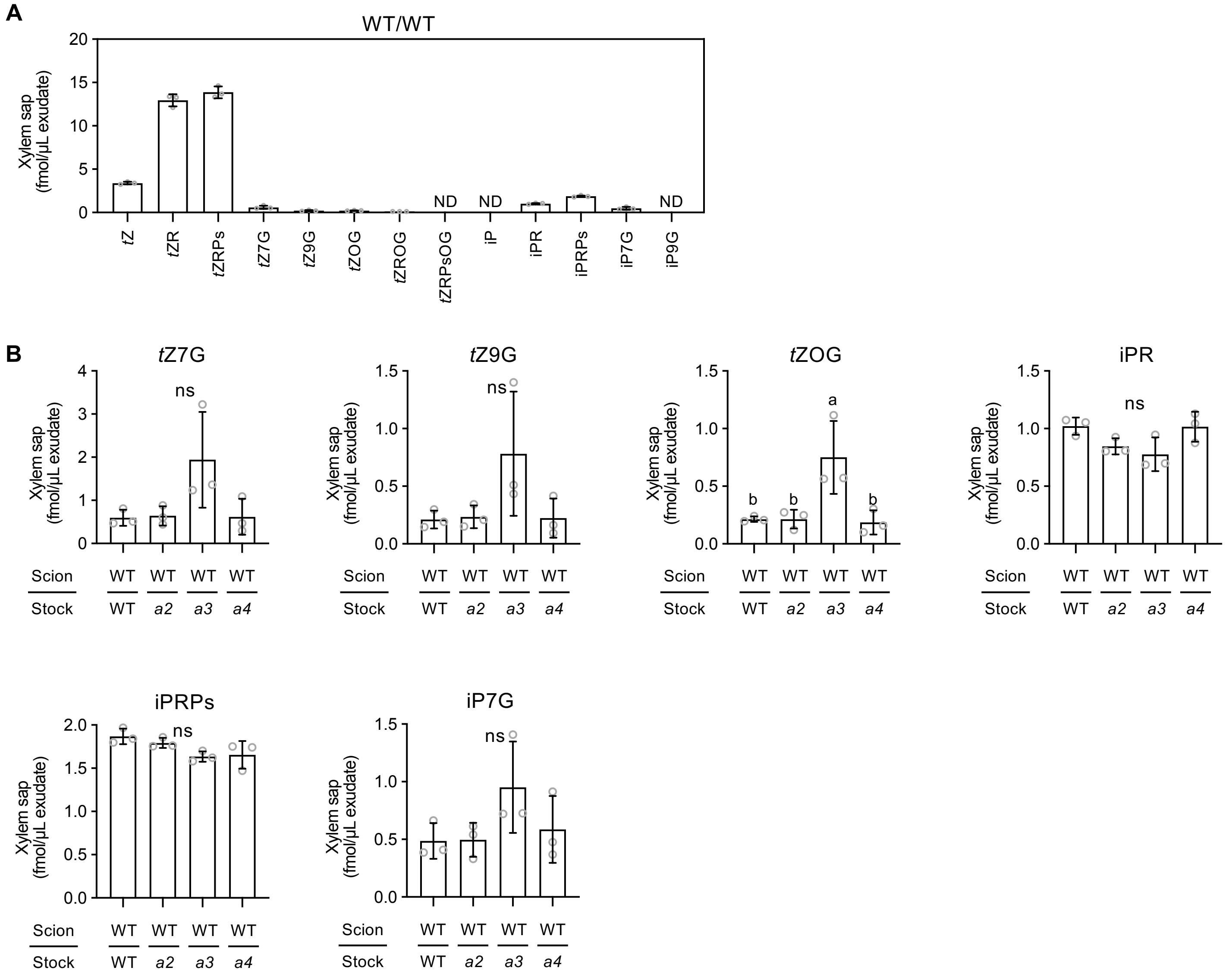


**Figure S7. Effects of root-specific CK receptor deficiency on xylem sap concentration of individual CK species.** (A) Xylem sap concentrations of *t*Z, *t*ZR, *t*ZRPs, *t*Z7G, *t*Z9G, *t*ZOG, *t*ZROG, iP, iPR, iPRPs, iP7G, and iP9G in WT/WT. (B) Xylem sap concentrations of *t*Z7G, *t*Z9G, *t*ZOG, iPR, iPRPs, and iP7G in grafted plants. Data pooled from three independent grafting experiments are presented as mean ± SD (n = 3). Different lowercase letters indicate significant differences, as determined via Tukey–Kramer test ($P < 0.05$). Grafted plants are denoted as "scion line/rootstock line." WT, Col-0; *a2*, *ahk2-5*; *a3*, *ahk3-7*; *a4*, *cre1-2; tZ*, *trans*-zeatin; *t*ZR, *t*Z-riboside; *t*ZRPs, *t*ZR 5′-phosphates; *t*Z7G, *t*Z-7 *N*-glucoside; *t*Z9G, *t*Z-9 *N*-glucoside; *t*ZOG, *t*Z-*O*-glucoside; *t*ZROG, *t*ZR-*O*-glucoside; *t*ZRPsOG, *t*ZRPs-*O*-glucoside; iP, $N^6$-($\Delta^2$-isopentenyl) adenine; iPR, iP-riboside; iPRPs, iPR 5′-phosphates; iP7G, iP-7 *N*-glucoside; iP9G, iP-9 *N*-glucoside; ND, not detected; ns, not significant.

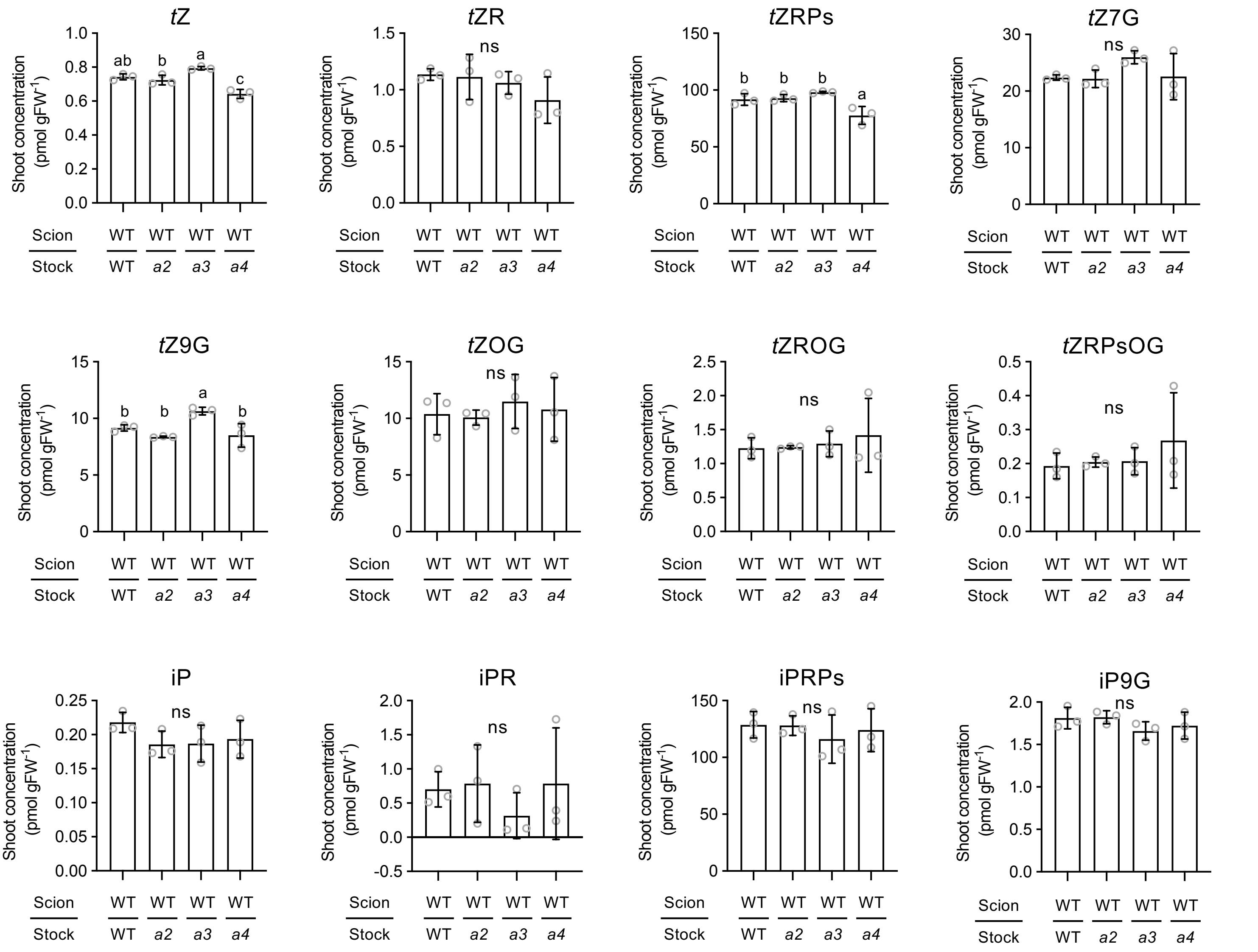


**Figure S8. Effects of root-specific CK receptor deficiency on shoot concentration of individual CK species.** Shoot concentrations of *t*Z, *t*ZR, *t*ZRPs, *t*Z7G, *t*Z9G, *t*ZOG, *t*ZROG, iP, iPR, iPRPs, and iP9G in grafted plants. Data pooled from three independent grafting experiments are presented as mean ± SD (n = 3). Different lowercase letters indicate significant differences, as determined via Tukey–Kramer test ($P < 0.05$). Grafted plants are denoted as "scion line/rootstock line." WT, Col-0; *a2*, *ahk2-5*; *a3*, *ahk3-7*; *a4*, *cre1-2;* FW, fresh weight; *t*Z, *trans*-zeatin; *t*ZR, *t*Z-riboside; *t*ZRPs, *t*ZR 5′-phosphates; *t*Z7G, *t*Z-7 *N*-glucoside; *t*Z9G, *t*Z-9 *N*-glucoside; *t*ZOG, *t*Z-*O*-glucoside; *t*ZROG, *t*ZR-*O*-glucoside; *t*ZRPsOG, *t*ZRPs-*O*-glucoside; iP, $N^6$-($\Delta^2$-isopentenyl) adenine; iPR, iP-riboside; iPRPs, iPR 5′-phosphates; iP9G, iP-9 *N*-glucoside; ns, not significant.

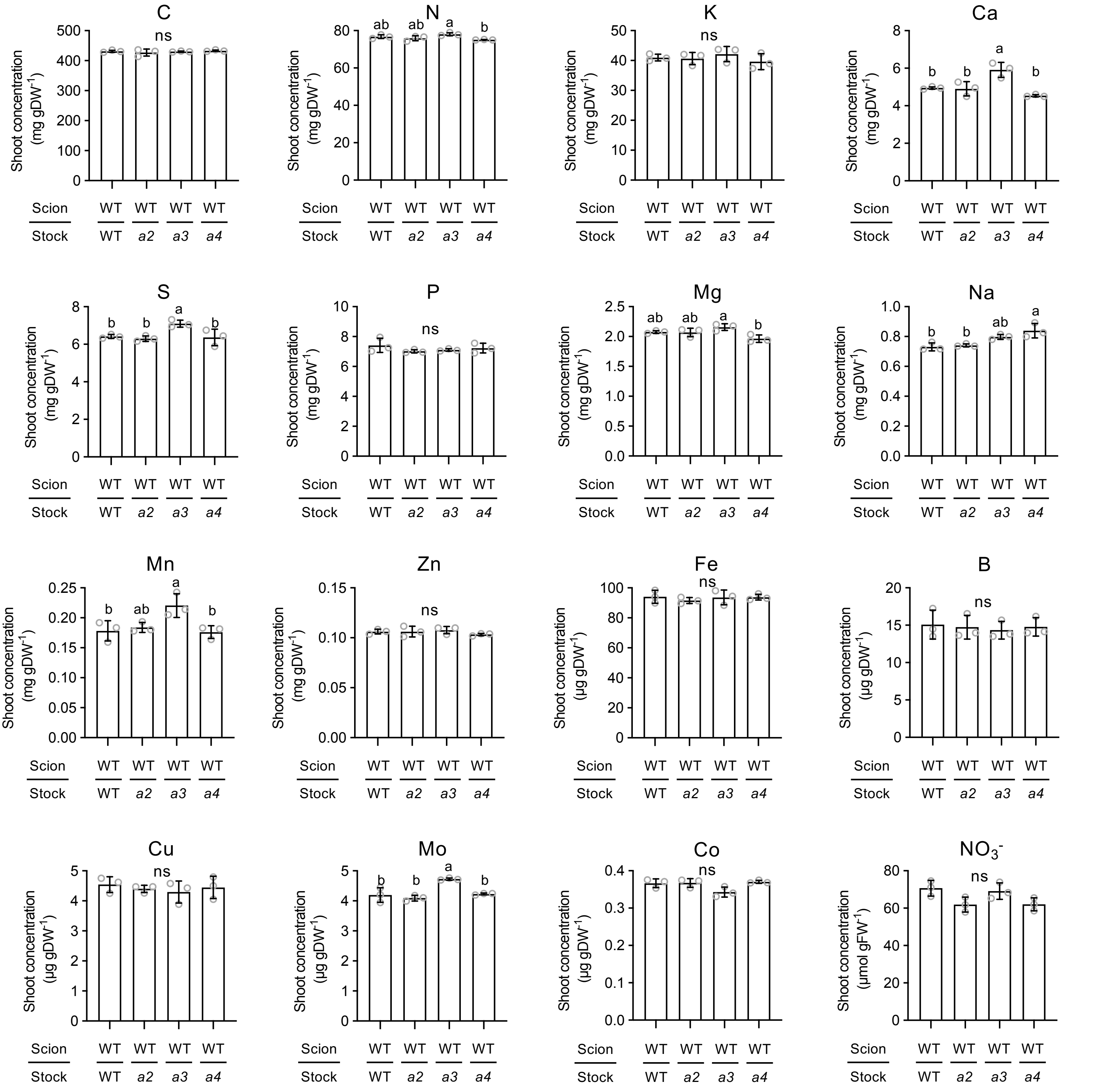


**Figure S9. Effects of root-specific CK receptor deficiency on shoot element concentrations.** Shoot concentrations of carbon and 14 nutrient elements and nitrate in grafted plants. All data pooled from three independent grafting experiments are presented as mean ± SD (n = 3). Different lowercase letters indicate significant differences, as determined via Tukey–Kramer test ($P < 0.05$). Grafted plants are denoted as "scion line/rootstock line." WT, Col-0; *a2*, *ahk2-5*; *a3*, *ahk3-7*; *a4*, *cre1-2;* DW, dry weight; FW, fresh weight; C, carbon; N, nitrogen; K, potassium; Ca, calcium; S, sulfur; P, phosphorus; Mg, magnesium; Na, sodium; Mn, manganese; Zn, zinc; Fe, iron; B, boron; Cu, copper; Mo, molybdenum; Co, cobalt; $NO_3^-$, nitrate; ns, not significant.

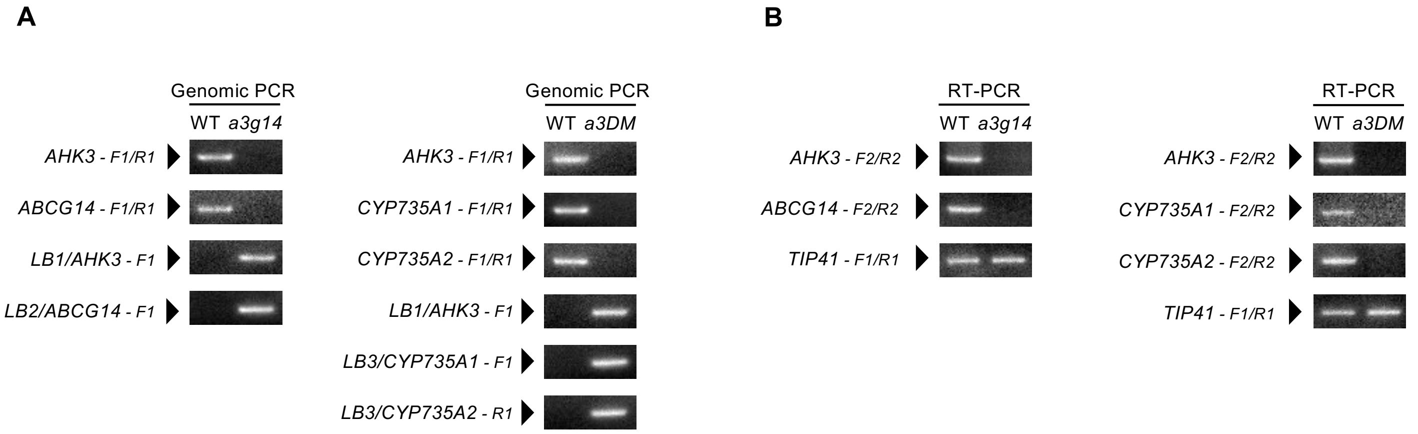


**Figure S10. Genomic PCR and RT-PCR results for *ahk3 abcg14* and *ahk3 cypDM*.** (A) Genomic PCR analysis of *AHK3*, *ABCG14*, *CYP735A1*, *CYP735A2*, and T-DNA left border in WT, *ahk3 abcg14*, and *ahk3 cypDM* seedlings grown 5 days on 1/2 MS medium. (B) RT-PCR analysis of *AHK3*, *ABCG14*, *CYP735A1*, and *CYP735A2* in WT, *ahk3 abcg14*, and *ahk3 cypDM* seedlings grown 7 days on 1/2 MS medium. *TIP41* was the control. Primers are listed in Supplementary Table S8. WT, Col-0; *a3g14*, *ahk3-7 abcg14*; *a3DM*, *ahk3-7 cyp735a1-2 cyp735a2-2*; *LB*, left border.

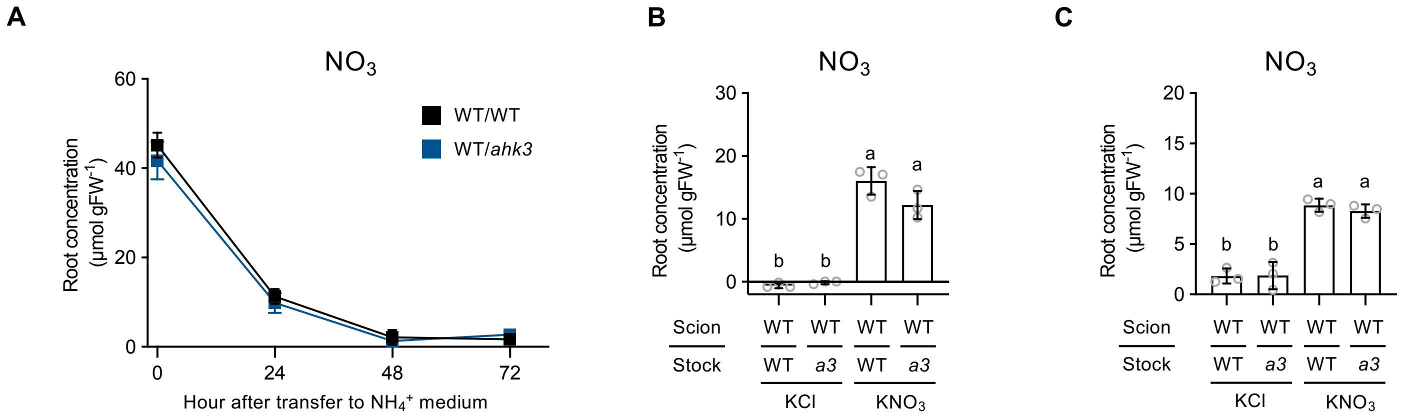


**Figure S11. Effects of root-specific *AHK3* deficiency on root nitrate concentration in response to nitrate signaling.** (A–C) Root nitrate concentration was measured at different time points and treatment conditions: (A) time course from 0 to 72 hours after transfer from 1/2 MS medium to 5 mM $NH_4^+$; (B) at 5 days or (C) 4 hours after transfer from 5 mM $NH_4^+$ medium to either 5 mM $NH_4^+$ + 0.5 mM $KNO_3$ or 5 mM $NH_4^+$ + 0.5 mM KCl. Data are presented as mean ± SD (n = 3). Different lowercase letters indicate significant differences, as determined via Tukey–Kramer test ($P < 0.05$). Grafted plants are denoted as "scion line/rootstock line." WT, Col-0; *a3*, *ahk3-7*; ns, not significant; FW, fresh weight; $NO_3$, nitrate; KCl, 5 mM $NH_4^+$ + 0.5 mM KCl; $KNO_3$, 5 mM $NH_4^+$ + 0.5 mM $KNO_3$.

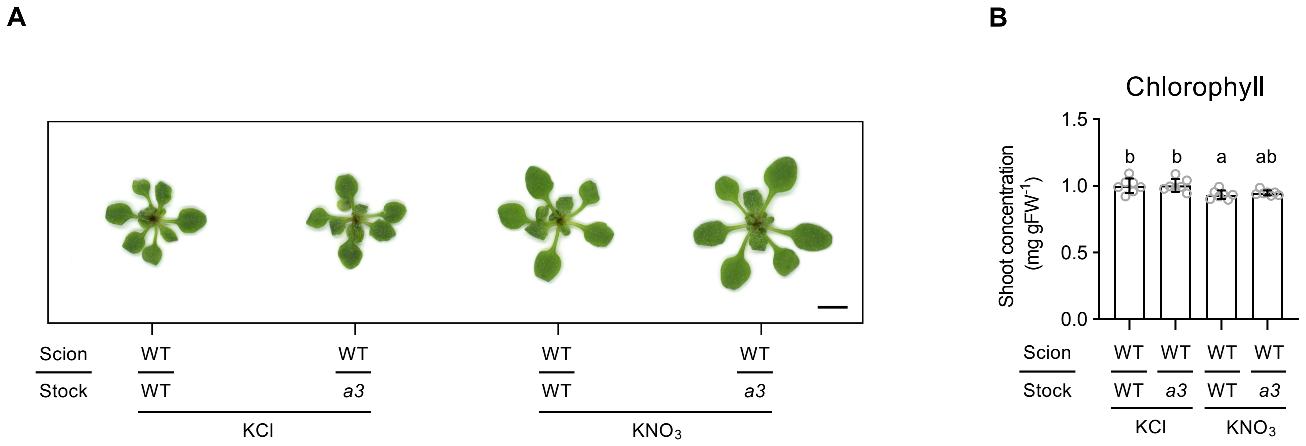


**Figure S12. Effects of root-specific *AHK3* deficiency on shoot chlorophyll concentration in response to nitrate signaling.** (A) Representative shoots of grafted plants 5 days after transfer from 5 mM $NH_4^+$ medium to either 5 mM $NH_4^+$ + 0.5 mM $KNO_3$ or 5 mM $NH_4^+$ + 0.5 mM KCl. Scale bars = 5 mm. (B) Shoot concentration of chlorophyll (a + b) in grafted plants at 5 days after transfer. Data pooled from two independent grafting experiments are presented as mean ± SD (n = 7). Different lowercase letters indicate significant differences, as determined via Tukey–Kramer test ($P < 0.05$). Grafted plants are denoted as "scion line/rootstock line." WT, Col-0; *a3*, *ahk3-7*; ns, not significant; FW, fresh weight; KCl, 5 mM $NH_4^+$ + 0.5 mM KCl; $KNO_3$, 5 mM $NH_4^+$ + 0.5 mM $KNO_3$.

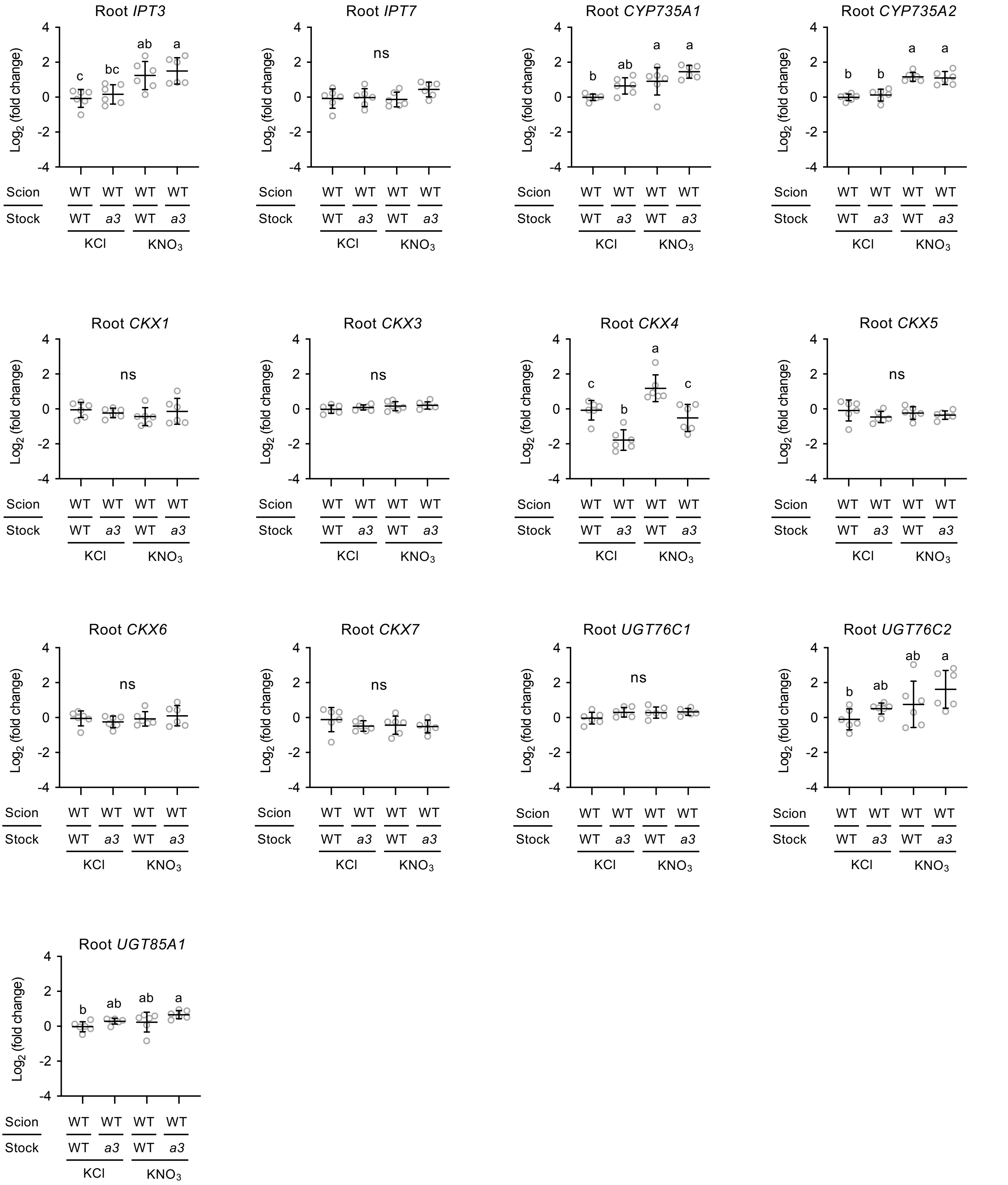


**Figure S13. Effects of root-specific *AHK3* deficiency on root gene expressions in response to nitrate signaling.** Root expression of *IPT3*, *IPT7*, *CYP735A1*, *CYP735A2, CKX1*, *CKX3*, *CKX4*, *CKX5*, *CKX6*, *CKX7*, *UGT76C1*, *UGT76C2*, and *UGT85A1* 4 hours after transfer from 5 mM $NH_4^+$ medium to either 5 mM $NH_4^+$ + 0.5 mM $KNO_3$ or 5 mM $NH_4^+$ + 0.5 mM KCl. Data pooled from two independent grafting experiments are presented as mean ± SD (n = 6). Graphs were generated from $\log_2$ fold change values relative to WT/WT under 5 mM $NH_4^+$ + 0.5 mM KCl (set = 1). Different lowercase letters indicate significant differences, as determined via Tukey–Kramer test ($P < 0.05$). Grafted plants are denoted as "scion line/rootstock line." WT, Col-0; *a3*, *ahk3-7*; ns, not significant; KCl, 5 mM $NH_4^+$ + 0.5 mM KCl; $KNO_3$, 5 mM $NH_4^+$ + 0.5 mM $KNO_3$.

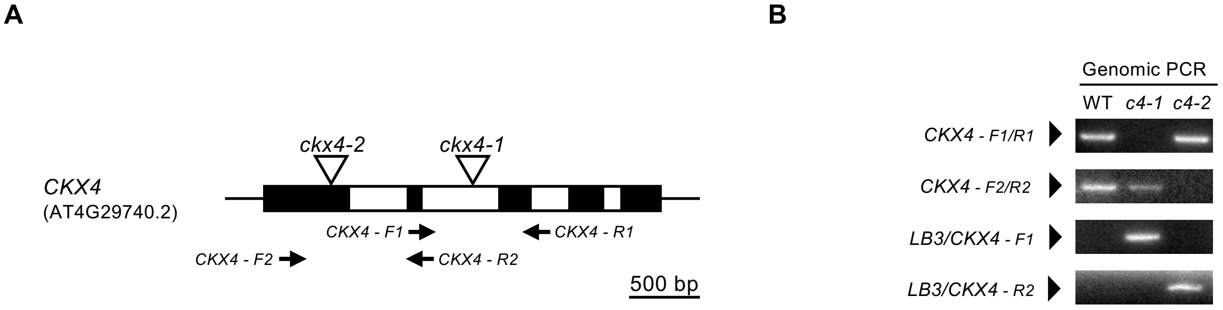


**Figure S14. Characterization of *CKX4* T-DNA insertion alleles.** (A) Schematic representative gene models for *CKX4* (AT4G29740.2) along with the positions of the primers used for genomic PCR. Site of T-DNA insertion in *ckx4-2* was determined by DNA sequencing using primers specific for the T-DNA left border. Black boxes represent exons, white boxes represent introns, and triangles indicate T-DNA insertion sites. Scale bar = 500 bases. (B) Genomic PCR analysis of *AHK3*, *CKX4*, and T-DNA left border in WT, *ckx4-1*, and *ckx4-2* seedlings grown 5 days on 1/2 MS medium. Primers are listed in Supplementary Table S8. WT, Col-0; *c4-1*, *ckx4-1*; *c4-2*, *ckx4-2*; *LB*, left border.

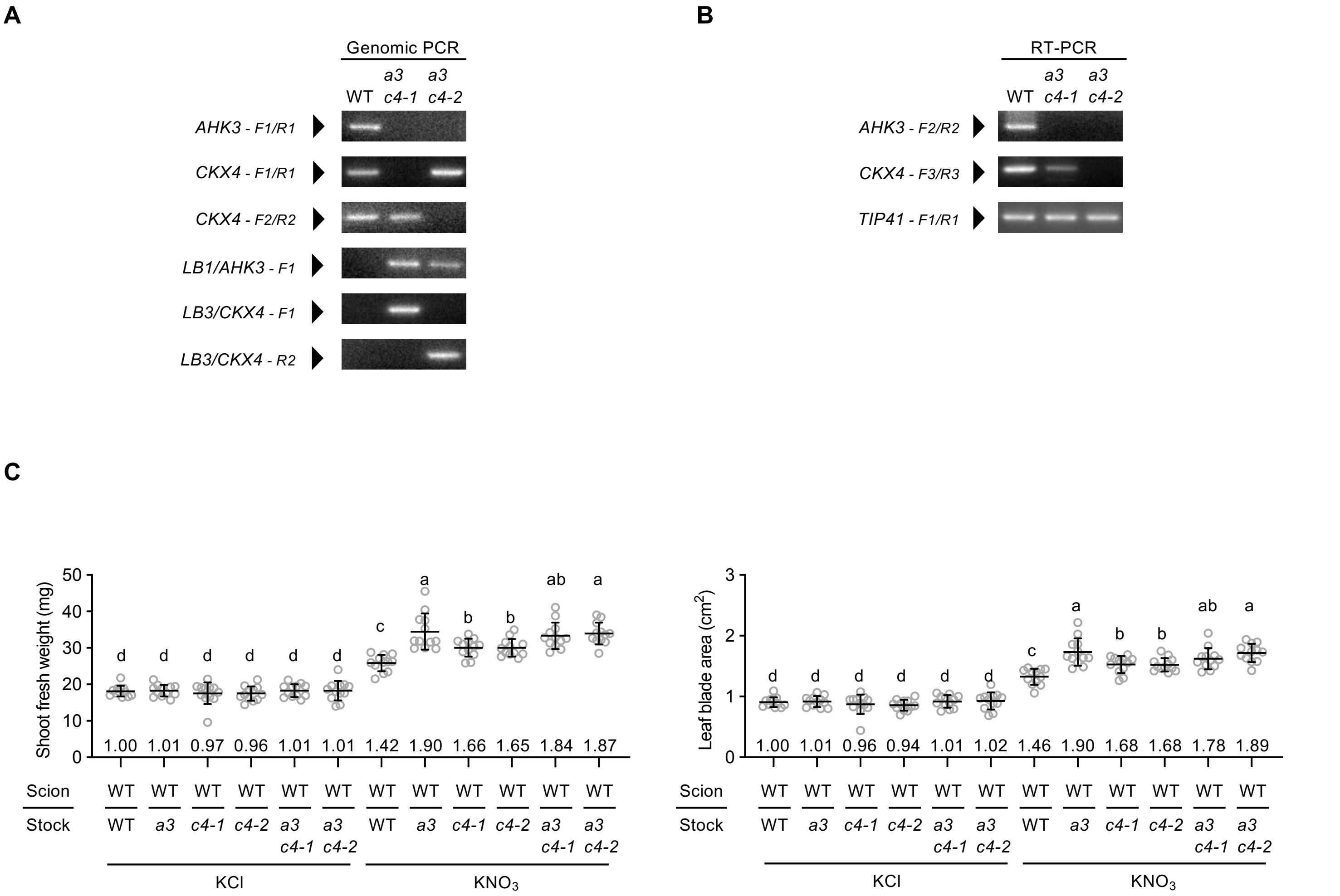


**Figure S15. Effects of root-specific deficiencies of *AHK3* and *CKX4* on shoot growth in response to nitrate signaling.** (A) Genomic PCR analysis of *AHK3*, *CKX4*, and T-DNA left border in WT, *ahk3 ckx4-1*, and *ahk3 ckx4-2* seedlings grown 5 days on 1/2 MS medium. (B) RT-PCR analysis of *AHK3* and *CKX4* in WT, *ahk3 ckx4-1*, and *ahk3 ckx4-2* seedlings grown 7 days on 1/2 MS medium. *TIP41* was the control. Primers are listed in Supplementary Table S8. (C) Shoot fresh weight and leaf blade area of WT/WT, WT/*ahk3*, WT/*ckx4-1*, WT/*ckx4-2*, WT/*ahk3 ckx4-1*, and WT/*ahk3 ckx4-2* 5 days after transfer from 5 mM $NH_4^+$ medium to either 5 mM $NH_4^+$ + 0.5 mM $KNO_3$ or 5 mM $NH_4^+$ + 0.5 mM KCl. Data pooled from three independent grafting experiments are presented as mean ± SD (n = 12). Different lowercase letters indicate significant differences, as determined via Tukey–Kramer test ($P < 0.05$). Graph numbers indicate mean values relative to WT/WT under 5 mM $NH_4^+$ + 0.5 mM KCl (set = 1). Grafted plants are denoted as “scion line/rootstock line.” WT, Col-0; *a3*, *ahk3-7*; *c4-1*, *ckx4-1*; *c4-2*, *ckx4-2*; *a3c4-1*, *ahk3-7 ckx4-1*; *a3c4-2*, *ahk3-7 ckx4-2*; KCl, 5 mM $NH_4^+$ + 0.5 mM KCl; $KNO_3$, 5 mM $NH_4^+$ + 0.5 mM $KNO_3$.

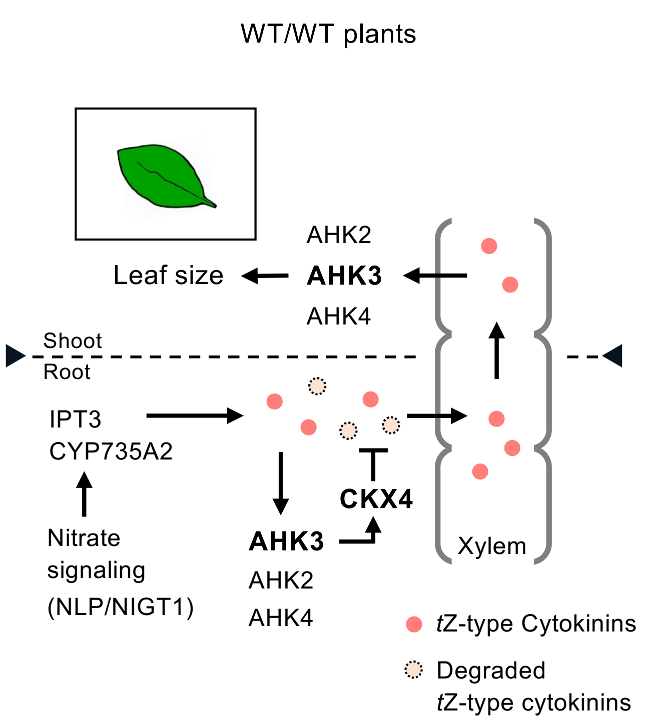


**Figure S16. A proposed model for xylem *tZ*-type CK transport controlled by AHK3 in response to nitrate signaling.** Root-expressed AHK3 modulates the xylem *tZ*-type CK transport, thereby controlling CK response and growth in leaves. The thickness of the lines represents the level of induction.

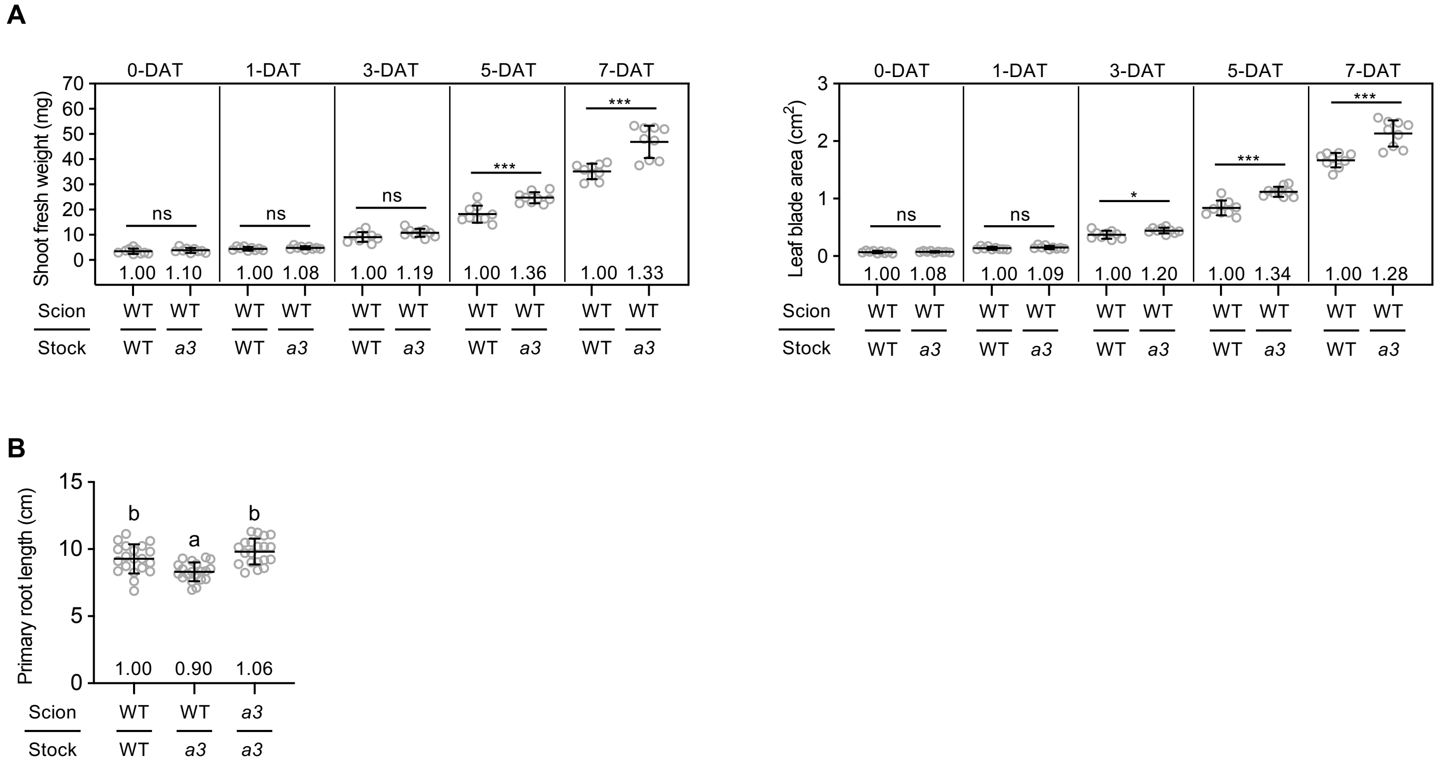


**Figure S17. Effects of root-specific *AHK3* deficiency on shoot and root growth.** (A) Time course of shoot fresh weight and leaf blade area in grafted plants after transfer to 1/2 MS medium. Data pooled from two independent grafting experiments are shown as mean ± SD (n = 9). Asterisks indicate significant differences (**P* < 0.05, ****P* < 0.001, Welch's *t*-test). (B) Primary root length of grafted plants 5 days after transfer to 1/2 MS medium. Data pooled from two independent grafting experiments are shown as mean ± SD (n = 20). Different lowercase letters indicate significant differences, as determined via Tukey–Kramer test (*P* < 0.05). Numbers on graphs show mean values relative to WT/WT (set = 1). Grafted plants are denoted as "scion line/rootstock line." WT, Col-0; *a3*, *ahk3-7*; ns, not significant; DAT, days after transfer.
